# Molecular Mechanisms Underlying Phenotypic Degeneration in *Cordyceps militaris*: Insights from Transcriptome Reanalysis and Osmotic Stress Studies

**DOI:** 10.1101/2023.08.29.555252

**Authors:** Chinh Q. Hoang, Giang H Dương, Mai H. Trần, Tao X. Vu, Tram B. Tran, T. N. Phạm Hằng

**Affiliations:** Center of Experimental Biology, National Center for Technical Progress, C6 Thanh Xuan Bac, Thanh Xuan, Hanoi, Vietnam; Center for Biomedical Informatics, Vingroup Big Data Institute, 458 Minh Khai, Hai Ba Trung, Hanoi, Vietnam; GeneStory JSC, 458 Minh Khai, Hai Ba Trung, Hanoi, Vietnam; Department of Pharmacology and Biochemistry, National Institute of Medicinal Materials, 3B Quang Trung, Hoan Kiem, Hanoi, Vietnam; University of Medicine and Pharmacy, Vietnam National University, 144 Xuan Thuy, Cau Giay, Hanoi, Vietnam

**Keywords:** Phenotypic degeneration, fungi, Cordyceps militaris, bioactive compounds, medicinal fungus, fruiting bodies, transcriptome reanalysis, MAPK signaling pathway, sexual reproduction, asexual sporulation, glycerol synthesis, osmotic stress, hyperosmotic stress, circles of areas, *Ste20*, Ste12, Gdp, Gpp, Gcy1, BrlA, *AbaA*

## Abstract

Phenotypic degeneration is a well-known phenomenon in fungi, yet the underlying mechanisms remain poorly understood. *Cordyceps militaris*, a valuable medicinal fungus with therapeutic potential and known bioactive compounds, is vulnerable to degeneration, which is a concern for producers. However, the causes of this process are still unclear. To shed light on the molecular mechanisms responsible for phenotypic degeneration in *C. militaris*, we isolated two strains with different abilities to form fruiting bodies. Our observations revealed that the degenerated strain had reduced ability to develop fruiting bodies, limited radial expansion, and increased spore density. We also conducted a transcriptome reanalysis and identified dysregulation of genes involved in the MAPK signaling pathway in the degenerate strain. Our RT-qPCR results showed lower expression of genes associated with sexual development and upregulation of genes linked to asexual sporulation in the degenerate strain compared to the wild-type strain. We also found dysregulation of genes involved in glycerol synthesis and MAPK regulation. Additionally, we discovered that osmotic stress reduced radial growth but increased conidia sporulation and glycerol accumulation in both strains, and hyperosmotic stress inhibited fruiting body formation in all neutralized strains. These findings suggest that the MAPK signaling pathway is dysregulated in the degenerate strain and the high-osmolarity glycerol and spore formation modules may be continuously activated, while the pheromone response and filamentous growth cascades may be downregulated. Overall, our study provides valuable insights into the mechanisms underlying *C. militaris* degeneration and identifies potential targets for future studies aimed at improving cultivation practices.

## Introduction

Phenotypic switching, also known as phenotypic degeneration, is a well-known occurrence in fungi. When cultivated in artificial conditions or subcultured repeatedly on artificial media, fungi undergo morphological changes such as growth retardation, altered pigmentation, decreased virulence, and reduced production of secondary metabolites, all of which result in batch-to-batch variations and negatively affect the quality and quantity of fungus-based products, making degenerate cultures a major concern for manufacturers (1). While genetic and/or epigenetic differences can contribute to degeneration, with some strains being more susceptible than others and certain gene mutations linked to phenotypic instability (2)(3)(4), the cultivation media can also play a role in this phenomenon. For example, cultures that are high in nutrients can accelerate degenerative phenotypes (5).

Fungi, like all living organisms, have the ability to sense and respond to changes in their environment and the MAPK pathway is one of the key signaling pathways that fungi use to respond to environmental cues. The MAPK pathway is an evolutionarily conserved signaling pathway found in all eukaryotic cells. It consists of a series of protein kinases that are activated in response to environmental signals. These kinases then phosphorylate downstream targets, including transcription factors, which ultimately modulate gene expression to adapt to a new condition (6). Fungi use the MAPK pathway for various processes such as cell cycle control, reproduction, morphogenesis, stress responses, cell wall assembly and integrity, virulence, and immunity (7). In the yeast *S. cerevisiae*, there are five branches of the MAPK pathway that mediate responses to different intrinsic and extrinsic clues, including response to pheromones during mating (pheromone response - PR), filamentous growth under nitrogen starvation conditions (filamentous growth - FG), response to osmotic stress and other stress conditions (high-osmolarity glycerol - HOG), response to cell wall stress (cell wall integrity - CWI), and spore formation (SF) (8). Mutations in two components of the MAPK pathway, *PaASK1* and *CpBck1*, have been linked to phenotypic instability in *Podospora anserine* (9) and to sectorization in *Cryphonectria parasitica* (*10*), but the role of the MAPK pathway in phenotypic degeneration is still not well understood.

The MAPK pathways initiate signal transduction through G-Protein-Couple Receptors, which detect intrinsic and extrinsic signals and transfer them to MAPK mediators such as CDC42 and STE20 for the PR, FG, and HOG modules, RHO1 for the CWI cascade, and unknown factor(s) for the SF branch. These MAPK regulators then activate MAPK kinase kinases such as STE11, which in turn phosphorylate MAPK kinases to activate MAPKs such as HOG1 and FUS3. The MAPKs then translocate to the nucleus and modulate the activity of transcription factors such as STE12 in the PR and FG modules, or HOT1 in the HOG cascade, which initiate transcription programs to adapt to new environments (10).

The PR, FG, and HOG modules share components but mediate distinct responses to different extracellular stimuli, indicating the existence of cross-pathway regulatory mechanisms. For example, both the FG and PR branches use the transcription factor STE12 to initiate transcription responses, but the STE12 and TEC1 heterodimer specifically induces genes resulting in filamentous growth (10). When responding to pheromone peptides, the MAPK FUS3 is activated and translocated to the nucleus, which simultaneously activates STE12 but also inactivates TEC1 to prevent the expression of genes for filamentous growth (11)(12). Similarly, the activation of the HOG cascade by hyperosmotic stress inhibits TEC1 activity and fine-tunes the signal for the stress response pathway (13). The presence of Hog1 and Pbs2 genes, two of the HOG module members, is essential for preventing the PR cascade from being activated by osmotic stress (14). The activated HOG branch also partially inhibits the mating response (15), and delays and dampens the PR cascade (16).

*Cordyceps militaris* is an ascomycete fungus that possesses medicinal properties such as immune modulation, anti-inflammation, anticancer, antidiabetic, anti-stroke, and anti-cardiovascular diseases. It has been used in traditional medicine in Asia for centuries and gained popularity in Europe and America in recent decades (17)(18). To meet market demands, *C. militaris* is artificially cultivated, but strain degeneration during subculturing and preservation is a major concern for manufacturers. Degenerated cultures exhibit a loss or reduction in growth rates, fruiting-body formation, pigmentation, and/or secondary metabolite production (19). Degenerated strains of C. militaris are associated with genetic and epigenetic changes, including mutations in the 18S and mating-type regions of the fungus (20). The degenerate strain also shows significantly higher DNA methylation, with differentially methylated genes enriched in pathways such as pyruvate metabolism, glycerophospholipid metabolism, DNA replication, and N-glycan biosynthesis (21). Gene expression changes are also linked to degeneration, with over 2000 differentially expressed genes identified in degenerated strains, including genes involved in toxin biosynthesis, energy metabolism, DNA methylation, and chromosome remodeling (22). These findings suggest that environmental stressors may cause genetic and epigenetic changes leading to phenotypic degeneration in *C. militaris* strains. However, the specific genes and pathways responsible for the phenotypes of the degenerate strain remain unknown.

In this study, we reanalyzed previously published gene expression data to identify biological pathways involved in degeneration and examined the dysregulated genes associated with the phenotypes of our degenerate C. militaris strain using RT-qPCR. We then manipulated the candidate pathway to confirm its involvement in phenotypic degeneration. Our study aims to provide a better understanding of the molecular mechanisms underlying phenotypic degeneration in fungi and to identify potential targets for preventing or mitigating its effects.

## Results

### Identifying indicators of phenotypic degeneration in *C. militaris*

The loss or reduction of fruiting body formation is a morphological hallmark of the *C. militaris* degenerate strain, which takes approximately two months to manifest. As a consequence, we wanted to find more rapid and reliable indicators of degeneration that can be easily measured and quantified. In our hands, radical expansions and spore density can be easily measured after twelve days of culture. Therefore, we quantify these two characteristics as indicators of *C. militaris* phenotypic degeneration in future studies.

We isolated two variations of the fungus *C. militaris* from the same batch of cultivation and conidial origin. One of these isolates, named *Ywt* (**Fig 1A**), is capable of forming fruiting bodies, while the other, named *Ydga* (**Fig 1B**), cannot. To confirm the genetic identity and relationship between the two variants, we conducted a BLAST search and phylogenetic analysis using the nucleotide sequences of the internal transcribed spacer (ITS) region of nuclear ribosomal DNA **(Additional File 1: Table S1)**. Our analysis revealed that the ITS sequences of *Ywt* and *Ydga* were identical to those of most, if not all, *C. militaris* specimens deposited in the National Library of Medicine, National Center for Biotechnology Information (NCBI). The neighbor joining tree analysis further revealed that *Ywt* and *Ydga* were closely related to other *C. militaris* strains **(Additional File 1: Figure S1)**.

**Figure 1:**
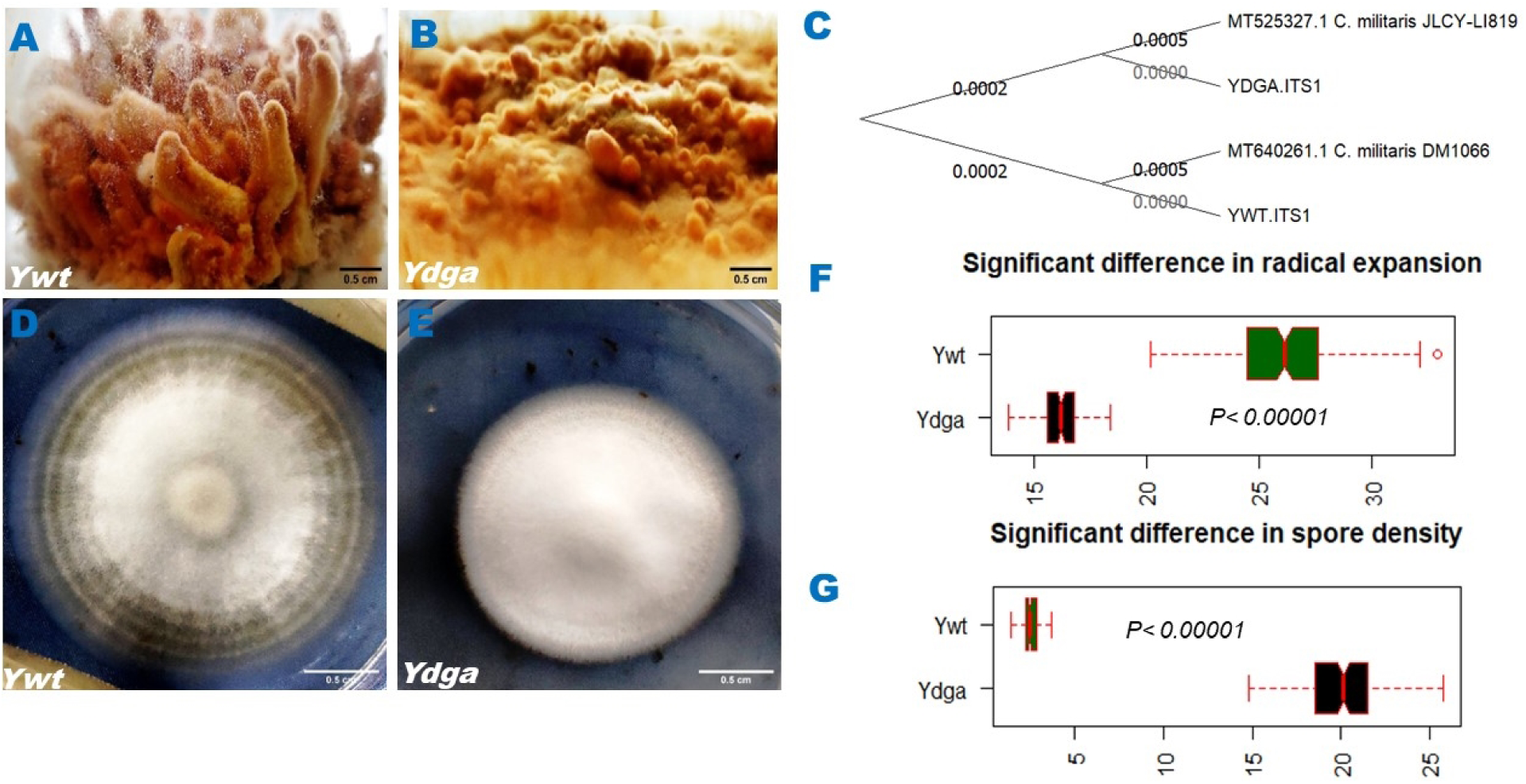
Characteristics of *C. militaris* phenotypic degeneration. (A vs. B) The neutralized strain (*Ywt*) developed fruiting bodies but the degenerate strain (*Ydga*) did not. (C) The phylogenetic tree of the *C. militaris* strains (interior-branch test with 1000 bootstrap replications). (D vs. E) Representative pictures of the two *C. militaris* isolates on PDA plates. (F & G) Significant differences (p < 0.00001) in areas of circles and spore densities of the two *C. militaris* variances. Data are presented as the means and standard deviations, with n = 5, and statistical significance was evaluated using a t test for two independent means.

To better understand the relationship between our *C. militaris* strains and other known strains, we compared them to the *DM1066* and *JLCY-LI819* strains through phylogenetic analysis. Our analysis showed that *Ydga* and *Ywt* were closely related to a common ancestor of the four strains, with a small interior branch length, while *DM1066* and *JLCY-LI819* were more distantly related (**Fig 1C)**. This suggests that *Ydga* and *Ywt* likely share an identical genetic background. Radical expansion reflects the growth rate and hyphal development of *C. militaris* strains on PDA plates. We found that *Ydga* had a significantly slower radical expansion than *Ywt*, indicating a lower growth rate and a defect in hyphal development. The hyphae of *Ydga* were irregular and fluffy, while those of *Ywt* were smooth and ring-shaped at the colony edge (**Fig 1D – 1F, Additional File 1: Table S2)**. These results suggest that the degenerate strain may have a reduced ability to adapt to the environment and utilize nutrients and/or may have halted genetic programs related to hyphal development.

Spore density reflects the asexual reproduction and sporulation of *C. militaris* strains. We found that *Ydga* had a significantly higher spore density than *Ywt*, indicating increased conidial formation (**Fig 1G, Additional File 1: Table S3**). This may be a compensatory mechanism for the loss of fruiting body formation, and/or a result of abnormal regulation of developmental pathways.

In conclusion, we have isolated and characterized two *C. militaris* variants with different abilities to form fruiting bodies. We have identified two morphological characteristics, radical expansion and spore density, that can be used as indicators of *C. militaris* strain degeneration. We have also revealed the phylogenetic relationship of our strains to other *C. militaris* strains based on ITS sequences.

### Transcriptome Analysis Identifies Dysregulated MAPK Signaling Pathway in Culture Degeneration of *C. militaris*

We performed transcriptome analysis to identify the molecular pathways involved in culture degeneration by comparing the gene expression profiles of a degenerate strain and a wild-type strain of *C. militaris*. We found that 880 genes were downregulated and 1034 genes were upregulated in the degenerate strain (a list of differentially expressed genes will be provided upon request). Among the downregulated genes, the most enriched gene ontology terms were ABC transporters, MAPK signaling pathway, and amino sugar and nucleotide sugar metabolism (**Fig. 2A, Additional File 2: Table S1)**, while no biological pathway was significantly enriched among the upregulated genes.

**Figure 2:**
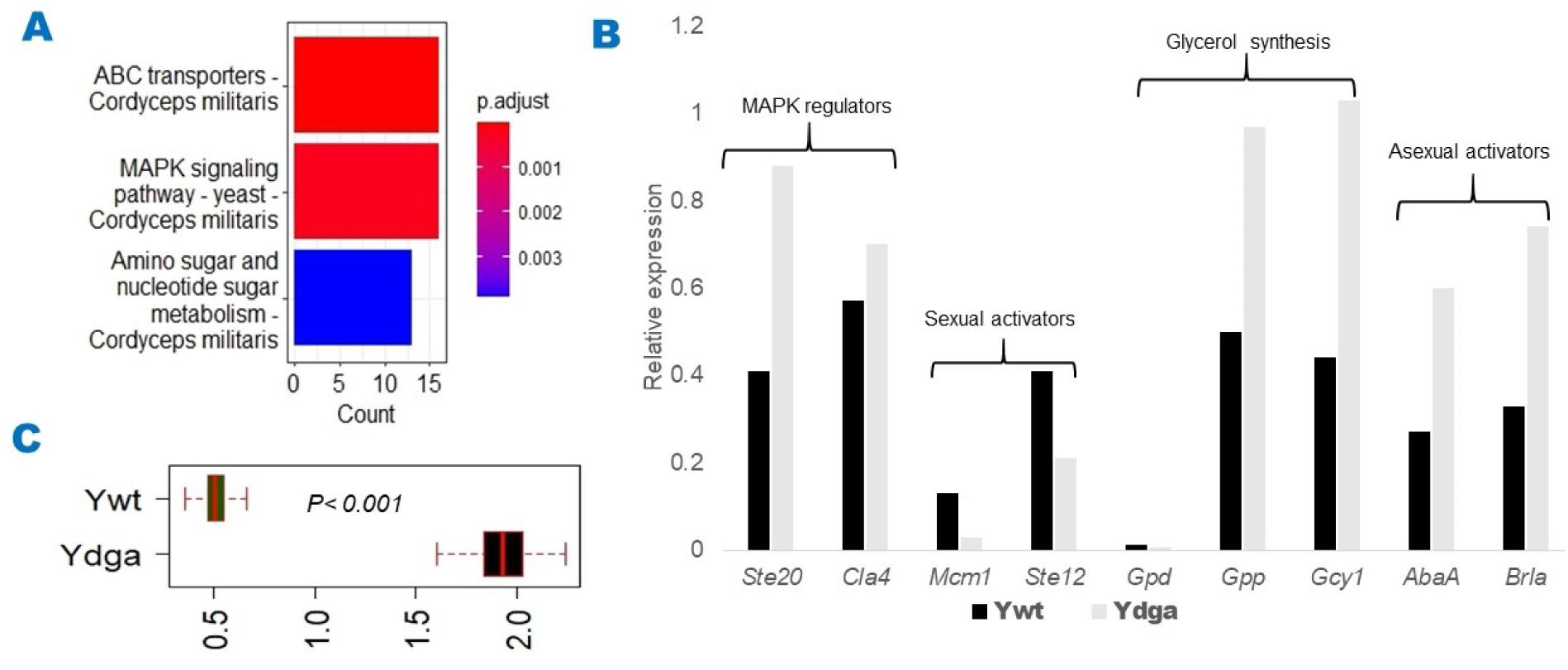
Dysregulation of the MAPK signaling pathway in *C. militaris* phenotypic degeneration. (A) Biological pathways significantly enriched in downregulated genes. (B) RT–qPCR results for genes involved in the MAPK pathway and associated with phenotypic degeneration i.e., genes contributing to sexual development, sporulation, and glycerol synthesis. (C) Intracellular glycerol contents of *Ywt* vs. *Ydga* were significantly different (p < 0.001). Data are presented as the means and standard deviations, with n = 5, and statistical significance was evaluated using a t test for two independent means.

We focused on the MAPK signaling pathway because it is responsible for regulating cellular processes such as sexual development and stress responses, which are affected by degeneration. To validate our findings, we used RT–qPCR to examine the expression levels of key genes involved in this pathway and associated with the phenotypes of our *C. militaris* degenerate strain, including genes related to sexual development (*Ste12, Mcm1*), asexual sporulation (*Brla, AbaA*), glycerol synthesis (*Gdh, Gpd*, and *Gpp*), and MAPK regulators (*Ste20, Cla4*) in the *Ywt vs. Ydga* strains. Our analysis found that the degenerate strain *Ydga* had lower expression of sexual development genes (*Ste12* and Mcm1) but higher expression of asexual sporulation genes (*Brla* and *AbaA*) than the wild-type strain *Ywt*. Additionally, the degenerate strain had higher expression of MAPK mediators (*Ste20* and *Cla4*) and glycerol-synthesizing enzymes (*Gdh* and *Gpp*) under hyperosmotic stress, but lower expression of the basal glycerol-synthesizing enzyme (*Gpd*) (**Fig 2B**). These findings were consistent with the intracellular glycerol content of the degenerate strain, which was elevated in comparison to the neutralized strain (**Fig 2C, Additional File 2. Table S2**). These results suggest that the MAPK signaling pathway is dysregulated in the degenerate strain, which may contribute to the observed phenotypic changes such as reduced radial growth, loss of fruiting body formation, and increased conidia sporulation. Overall, our study highlights the potential role of the MAPK signaling pathway in phenotypic degeneration.

### The HOG module may play a role in *C. militaris* phenotypic degeneration

Our observation of dysregulated transcription of MAPK signaling pathway components and elevated intracellular glycerol content in the degenerate strain of C. militaris led us to hypothesize that stress may contribute to the observed phenotypic degeneration, with the HOG module potentially playing a role. To test this hypothesis, we exposed both the wild - type strain *Ywt* and the degenerate strain *Ydga* to various stressors and measured the effects on radial growth, conidia sporulation, and glycerol accumulation.

We found that Congo red, a cell wall integrity (CWI) stressor, had similar effects on both strains, reducing radial growth and conidia sporulation to the same extent (**Fig 3A&B, Additional File 3. Table S1**). This suggests that the CWI branch is not involved in the phenotypic degeneration of *C. militaris*. However, we observed differential effects of oxidative and osmotic stressors on the two strains. H2O2, an oxidative stressor, slightly decreased radial growth but increased conidia sporulation in both strains, while N-acetyl cysteine (NAC), an antioxidant agent, inhibited radial growth and had no significant effects on conidia sporulation in the two strains (**Fig 3A & 3C, Additional File 3. Table S2**). These results suggest that oxidative stress may partially contribute to the phenotypic degeneration of *C. militaris*.

**Figure 3:**
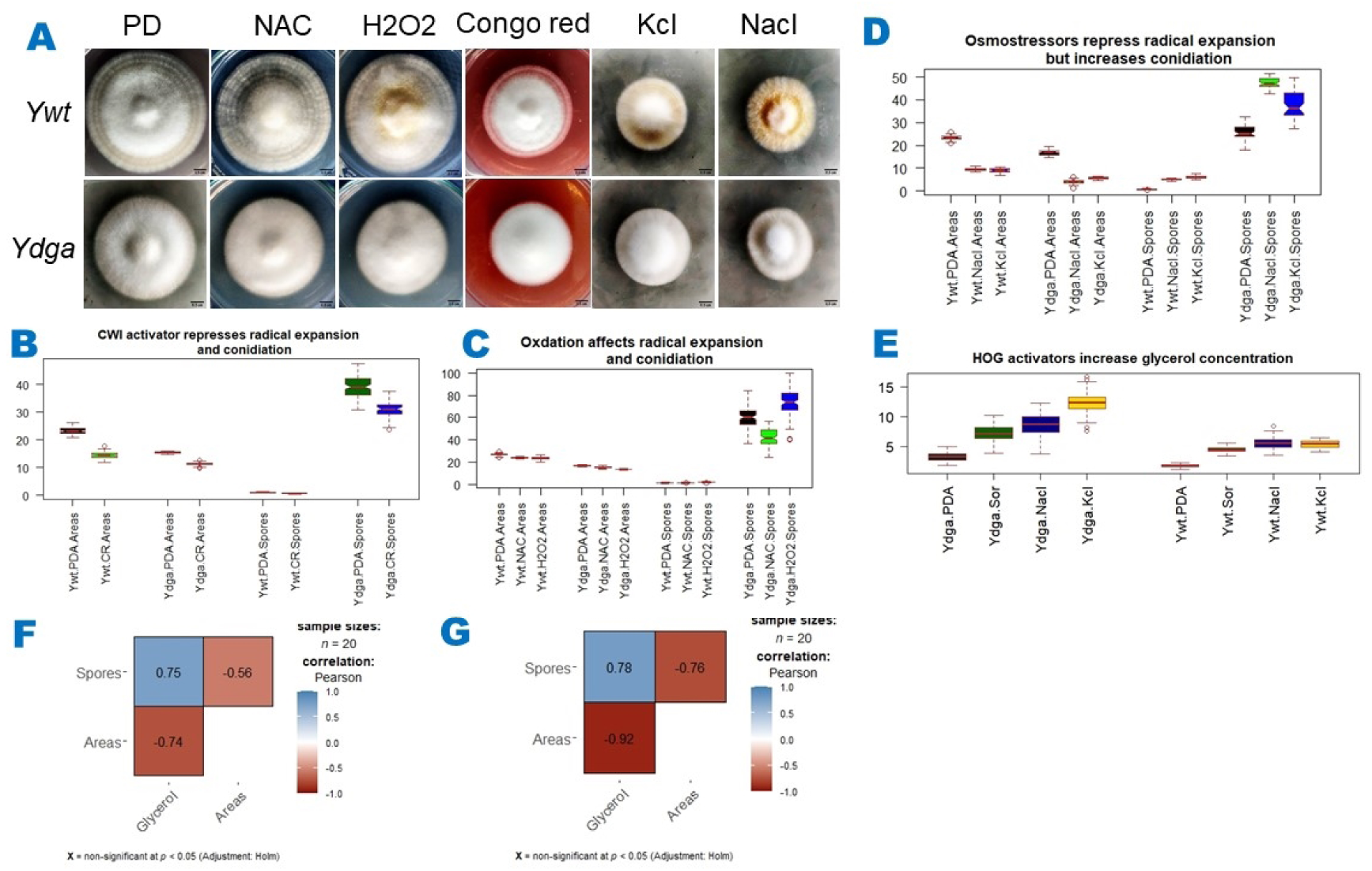
Effect of stressors on radical expansion, sporulation, and intracellular glycerol. (A) Representative images of *C. militaris Ywt* and *Ydga* cultured in PDA with or without different stressors. (B) Congo red, a CWI activator, significantly (p </= 0.006) repressed both radical expansion and sporulation in both C. militaris strains. (C) H2O2, an oxidative stressor, significantly (p < 0.05) suppressed radical expansion but promoted sporulation in both strains. NAC, an antioxidant agent, significantly (p < 0.05) inhibited radical growth without affecting sporulation in either strains. (D) KCl and NaCl, two osmotic stressors, significantly (p </= 0.002) suppressed radical expansion but promoted sporulation in both strains. (E) Osmotic stressors significantly (p < 0.00001) increased intracellular glycerol concentrations in both strains. (F & G) Glycerol concentrations showed a positive correlation with spore density but a negative correlation with circle areas in both strains. Data are presented as the means and standard deviations, with n = 5, and statistical significance was evaluated using a t test for two independent means.

The most notable effects on both strains were observed when exposed to osmotic stressors such as KCl (potassium chloride), NaCl (sodium chloride), or sorbitol. These stressors significantly reduced radial growth but increased conidia sporulation and glycerol accumulation in both strains (**Fig 3A, 3D & 3E, Additional File 3. Table S3, S4, S5**). Additionally, we observed that intracellular glycerol concentrations were positively and negatively correlated with spore density and radial expansion (**Fig 3F&G**), suggesting the existence of cross-regulatory mechanisms among the HOG, FG, and SP modules. Overall, our findings suggest that osmotic stress plays a significant role in the phenotypic degeneration of *C. militaris*, with the HOG cascade potentially being involved in this process.

### Hyperosmotic stress induces phenotypic degeneration in neutralized *C. militaris* strains

To investigate whether the involvement of the HOG cascade in *C. militaris* phenotypic degeneration is a widespread phenomenon, we studied the effects of hyperosmotic stress on four *C. militaris* strains, including three neutralized strains (*Nf, Wt*, and *Ywt)* and one degenerated strain (*Ydga*). We confirmed the origin and phylogenetic relationship of the *Nf* and *Wt* strains with the *Ywt, Ydga* and *JLCY-LI819* strains by ITS DNA sequencing. The resulting phylogenetic tree indicated that these strains share a common ancestor and that the neutralized strains (*Ywt, Wt* and *Nf*) belong to the same clade (**Fig 4B, Additional File 1. Table S1**).

**Figure 4:**
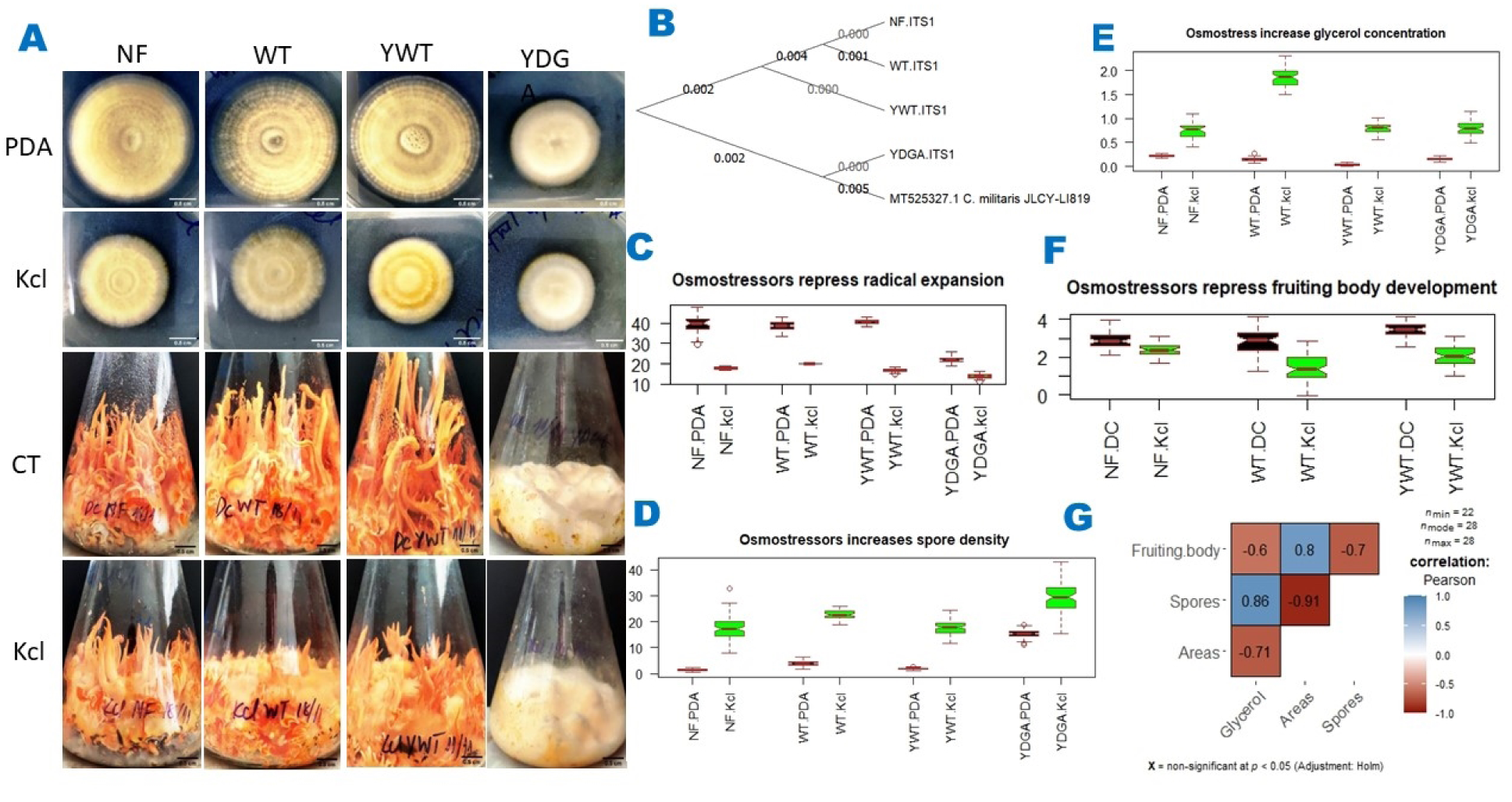
Hyperosmotic conditions induce phenotypic degeneration in neutralized *C. militaris* strains. (A) Representative images of radical (upper panel) and fruiting body (lower panel) repression by hyperosmotic cultures. (B) Phylogenetic tree. (C – F) Hyperosmotic conditions significantly (p </= 0.002) suppressed radical expansion and fruiting body development, but promoted spore densities and glycerol concentrations in all neutralized *C. militaris* strains. (G) Glycerol concentrations showed a positive correlation with spore density but a negative correlation with circle areas and fruiting body weights. PDA: Potatoes dextrose agarose; CT = FBM; Kcl = PDA or FBM + 0.4 M Kcl. Data are presented as the means and standard deviations, with n >/= 7, and statistical significance was evaluated using a t test for two independent means.

Our findings demonstrated that the addition of potassium chloride to PDA media decreased radial growth by approximately 70% in neutralized strains and 30% in degenerated strains (**Fig 4A-upper panel, 4C, Additional File 4. Table S1**). This implies that the degenerated strain may have a higher tolerance to osmotic stress compared to the neutralized strains, or that the degenerated strain has reached its maximum limit in terms of growth inhibition. Additionally, hyperosmotic stress increased spore production (**Fig 4D, Additional File 4. Table S1**) and glycerol accumulation (**Fig 4E, Additional File 4. Table S2**) in all strains. Interestingly, hyperosmotic stress inhibited fruiting body formation by approximately 50% in all neutralized strains, while the degenerated strain failed to produce fruiting bodies under any condition (**Fig 4A – lower panel, Fig 4F, Additional File 4. Table S3**).

Furthermore, we observed a positive correlation between glycerol concentration and spore density, and a negative correlation between glycerol concentration and radial growth area and fruiting body weight (**Fig 4G**). Taken together, our results suggest that in *C. militaris* degenerated strains, the HOG and SP modules may be continuously activated, while the PR and FG cascades may always be deactivated, contributing to the observed phenotypic degeneration.

## Discussion

*Cordyceps militaris* is an ascomycete mushroom with well-known nutritional and medicinal properties. However, its long-term cultivation poses a significant challenge due to the degeneration of strains, which affects its commercial production. The objective of this article is to investigate the molecular basis of phenotypic degeneration in *C. militaris.* Degenerate strains lose the ability to produce fruiting bodies, show reduced radical expansion, and increased conidia density. Transcriptome data analysis reveals dysregulation of several molecular pathways, including the MAPK signaling pathway, which regulates various cellular activities and has been linked to degeneration. RT–qPCR analysis showed altered expression levels of key genes involved in the MAPK pathway, which were associated with the observed phenotypes of our *C. militaris* degenerate strain. Our results also indicate that hyperosmotic stress induces phenotypic degeneration in *C. militaris* and that continuous activation of the HOG module may be a contributing factor to *C. militaris* phenotypic degeneration.

The degenerate strain of *C. militaris* is characterized by the loss or reduction of fruiting body formation, which has been used as an indicator for further studies (23)(24)(22). However, this phenotype takes approximately two months to manifest. Therefore, we aimed to identify morphological phenotypes that are easy to quantify and require a shorter time to manifest. We isolated two *C. militaris* variations from the same cultivated batch, one with the ability to produce fruiting bodies and the other without on solid media. We compared their radical expansion after 12 days of culture on PDA plates and found that the degenerate strain reduced radical expansion by approximately 50% compared to the wild type strain. While Wellhan et al. (2021) reported nonsignificantly slower radical growth rates in the degenerate strain, they did not show its ability to form fruiting bodies, so it is unclear whether the “degenerate” strain is truly degenerate (25). Our study revealed that the degenerated strain showed a significantly higher conidia density than the wild-type strain, which contradicts the findings of Meiyu et al. (2022).

Their study reported a significantly lower total number of conidia per dish in the degenerate strain by day 20 of culture (26). This discrepancy may be due to differences in the strains studied, sample collection times, and methodologies used.

We conducted a reanalysis of the transcriptome data generated by Yin et al. (2017) to identify the signaling pathways involved in the phenotypic degeneration of *C. militaris*. Our analysis revealed over 1000 differentially expressed genes, and the pathways of ABC transporters, MAPK signaling, and amino sugar and nucleotide sugar metabolism were significantly enriched among downregulated genes. In contrast, Yin et al. identified over 2000 differentially expressed genes including those involved in toxin biosynthesis, energy metabolism, DNA methylation, and chromosome remodeling (22). This discrepancy may have arisen from differences in the bioinformatics pipelines and/or the datasets used for analysis.

We focused on the MAPK signaling pathway because it regulates various cellular processes such as sexual development and stress responses, and its dysregulation has been associated with phenotypic switching in other fungal species (10)(7)(9)(27). To verify the expression levels of key genes involved in the MAPK pathway that are associated with the phenotypes of our C. militaris degenerate strain, we used RT–qPCR to compare the transcription levels of genes involved in sexual development, asexual sporulation, glycerol synthesis, and MAPK activators. The RT–qPCR results are consistent with the observed phenotypes of the *C. militaris* degenerate strain (*Ydga*), with downregulation of sexual development genes (*Ste12* and *Mcm1*) and upregulation of conidiation genes (*BrlA* and *AbaA*). Additionally, the expression of MAPK- activator (*Ste20*) and glycerol synthesis (*Gpp* and *Gdh*) genes was doubled in *Ydga* compared to the *Ywt* strain.

In our study, we found that the *C. militaris* ortholog transcripts of the yeast STE12 and MCM1 transcription factors were downregulated. STE12 is a downstream target of the PR module that regulates genes involved in sexual development in response to pheromone signals. Additionally, STE12 physically interacts with TEC1 to control genes involved in filamentous growth under starvation conditions and physically cooperates with MCM1 to mediate the expression of pheromone-inducible genes necessary for proper sexual development (28)(29)(30)(31).

However, the specific role of STE12 and STE12-like proteins in fruiting body formation varies between species (28). MCM1 function is required for fruiting body development in the homothallic ascomycete *Sordaria macrospora*, as well as growth, conidiogenesis, cell wall integrity, and the cell cycle in the filamentous insect-pathogenic fungus *Beauveria bassiana* (32)(33). These data indicate that the downregulation of the *Ste12*- and *Mcm1*- orthologous genes in the *C. militaris* degenerate strain *Ydga* may contribute to growth and fruiting body retardation.

The central regulatory pathway for conidiogenesis, involving three sequentially controlled transcription factors (BRLA, ABAA, and WETA), is conserved in Ascomycete fungi (34). In Aspergillus, BRLA is the essential activator of asexual sporulation, and its deletion results in failure to develop vesicle structures and instead produces elongated, bristle-like aerial stalks (35). The deletion of *brlA* also abolishes the production of other conidiation-specific genes such as *AbaA, WetA, VosA* and *RodA* (36). *AbaA* is required for the differentiation of phialides, and its loss of function results in the production of cylinder-like terminal cells with no conidia being formed. *WetA* functions in completing conidiogenesis, and its loss results in colorless conidia with defective spore walls (35). This regulatory network, which is regulated by the Slt2-MAPK/RNS1, Fus3-MAPK and Hog1-MAPK cascades, also regulates asexual development in *Metarhizium*, and their loss of function impairs conidiogenesis (37). These data suggest that upregulation of the BrlA and AbaA transcripts in the *C. militaris* degenerate strain *Ydga* may lead to an increase in its conidia density.

Glycerol synthesis and accumulation play a major role in the response to hyperosmotic and oxidative stresses (38). Yeast GPD is the rate-limiting enzyme for glycerol biosynthesis (39), while in *Aspergillus nidulans*, GLD (GCY1 orthologous) is crucial for adapting to hyperosmotic stress(40). In *Cryptococcus neoformans*, GPP2 is a key enzyme in response to various stresses (41). In our study, we found that the transcription levels of *Gcy1* and *Gpp* were upregulated, while that of *Gpd* was downregulated in the degenerate strain *Ydga*, indicating that the elevated glycerol content in the *Ydga* strain may partially depend on GLD and GPP but not on GPD. The *Ydga* strain also increased the expression of *Ste20* and *Cla4* transcripts, which are involved in glycerol biosynthesis enzymes in yeast.

The degenerated strain *Ydga* exhibited higher levels of glycerol concentration and glycerol biosynthesis enzymes, indicating that the HOG module may be persistently activated in the degenerate strain. In support of this notion, our experiments under hyperosmotic conditions revealed that all three neutralized *C. militaris* strains underwent phenotypic degeneration, characterized by suppressed fruiting body and hyphal development, but increased conidia density and glycerol accumulation. Moreover, the radical growth of *C. militaris* and the expression of the *Fus3* and *Hog1* genes were repressed in a concentration-dependent manner under hyperosmotic conditions (42). Taken together, these findings suggest that the continuous activation of the HOG cascade under artificial culture conditions may lead to the dysregulation of MAPK pathway genes and phenotypic switching.

Based on our data, the degenerate strain *Ydga* exhibited higher levels of Ste20 transcripts, a shared activator of the PR, FG, and HOG cascades, but lower expression of the Ste12 transcript, a crucial transcription factor of the PR and FG branches. Consequently, the HOG cascade may be persistently activated due to the continuous upregulation of *Ste20* transcripts, while the PR and FG cascades may be downregulated owing to the decrease in Ste12 transcripts, leading to defects in fruiting body development, a reduction in radical growth, and an increase in conidia density.

Our model provides a useful framework to investigate the molecular mechanisms that underlie the contribution of environmental stress to phenotypic degeneration in fungi. For instance, it raises questions about whether stress alters transcripts at the genetic, epigenetic, or RNA metabolism level, or whether overexpression of *Ste20/Cla4* in a *Ste12* inactivation background results in phenotypic degeneration. These questions could be explored in future studies to gain a better understanding of the molecular mechanisms that govern phenotypic degeneration in fungi.

This study represents the first to demonstrate that radical growth and conidia density can be used as rapid indicators of *C. militaris* phenotypic degeneration, that the MAPK signaling pathway is transcriptionally dysregulated and intracellular glycerol content is elevated in the degenerate strain, and that stress, such as hyperosmotic stress, may contribute to phenotypic degeneration in *C. militaris.* Additionally, the study suggests that the HOG module may be continuously activated in the degenerate strain. The findings also open up new research avenues on the role of the stress response and the MAPK pathway in fungal physiology and morphology, including the relationship between glycerol metabolism and fungal phenotypic plasticity. The implications of this work could extend beyond *C. militaris* to other filamentous fungi, with potential applications in enhancing the yield of bioactive compounds and understanding the regulation of fungal morphology and physiology. The results emphasize the importance of understanding the molecular mechanisms underlying degeneration in filamentous fungi, as this knowledge could aid in preventing degeneration during industrial production and developing early detection and monitoring tools for degenerative strains. Overall, this study provides significant insights into the molecular basis of phenotypic degeneration in *C. militaris* and highlights its potential impact on industrial cultivation.

Several limitations need to be considered in our presented findings. First, our study only focused on two morphological characteristics, radical expansion, and spore density, as indicators of phenotypic degeneration in *C. militaris*. While these indicators may be useful, they may not capture the full spectrum of changes that occur during degeneration. Therefore, further studies should investigate additional indicators and utilize multiple criteria to assess phenotypic changes. Second, our study only examines one *C. militaris* degenerated strain. As such, our findings may not be representative of the broader population of *C. militaris* strains. Therefore, caution should be exercised when generalizing the results to other strains. Third, our study relied on transcriptome analysis to identify molecular pathways involved in culture degeneration. While transcriptome analysis provides valuable insights into gene expression changes, it does not necessarily reflect protein expression or activity. Thus, it is important to use alternative approaches to confirm these results and conduct further investigations. Fourth, our study only examined the role of the MAPK signaling pathway in *C. militaris* phenotypic degeneration. Other molecular pathways may also be involved in this process, and further studies should explore their potential contributions. Finally, our study uses stressors to investigate the involvement of the HOG cascade in *C. militaris* phenotypic degeneration. While stressors may be useful in inducing and studying cellular responses, they may not accurately reflect the conditions under which degeneration occurs in natural environments. Therefore, further investigation using more ecologically relevant conditions is necessary to confirm the findings.

### Conclusions

Our study provides significant insights into the molecular basis of phenotypic degeneration in *C. militaris*. The identification of easily quantifiable indicators, the dysregulation of the MAPK signaling pathway, and the role of the stress response in phenotypic degeneration contribute to our understanding of this complex phenomenon. Future studies should further investigate additional indicators, examine a broader range of strains, confirm results through alternative approaches, and explore the involvement of other molecular pathways. Ultimately, these findings will aid in preventing degeneration during cultivation and advancing the field of fungal biology.

## Methods

### Fungal Strain and Media

Strains of neutralized *C. militaris* (*Nf* and *Wt*) were gifted by Vu Duy Nhan, while strains of neutralized (*Ywt*) and degenerate (*Ydga*) *C. militaris* were gifted by Le Thi Hai Yen from the Laboratory of Macro Fungi Technology, Institute of Microbiology and Biotechnology. The strains were cultured on potato dextrose agar (PDA) media containing 200 g/L potato, 20 g/L glucose, and 2 g/L agar, or potato dextrose broth (PDB) media containing 200 g/L potato and 20 g/L glucose. Fruiting body media (FBM) was prepared by mixing 35 grams of brown rice with 70 mL of liquid media containing 200 g/L potato, 20 g/L glucose, and 100 g/L silkworm pupae.

#### Phenotypic Analysis

To measure colony size and conidia density, the *C. militaris* strains were cultured on PDA media. Ten microliters of 10^6^ conidia were dropped in the center of a PDA Petri dish and inoculated for 12 days under natural daily dark-light cycles. The diameters of the colonies were measured and converted to circles of areas using the formula {(diameter/2)^2^ x 3.14}. The conidia were harvested in the sterile 0.01% Triton X-100 (MERK, cat# 9036-19-5) and filtered through milk filters (Lamtor Ltd, Bolton, UK) to remove hyphal fragments. The conidia were counted with a hemocytometer and their density was calculated using the formula [(conidia concentration x number of mL of the filtered suspension)/circles of areas of corresponding colony].

To measure fruiting body weight, the *C. militaris* conidia were cultured in PDB media at 130 rpm and 25°C for 3 days. The inoculated media were then added to the FBM and incubated in the dark for 2 weeks, followed by 6 weeks in a 12/12 dark-light cycle with 90% humidity. The fruiting bodies were collected, dried at 80°C, and weighed. To assess hyperosmotic stress, the *C. militaris* strains were cultured on PDA, PDB, or FBM supplemented with 0.4 M KCl, 0.4 M NaCl, or 1 M sorbitol.

#### Glycerol Measurement

The *C. militaris* hyphae were collected from 6-day-old cultures on PDA plates, and their weight was recorded. Then, 1.2 mL of 50% EtOH was added to each tube, and the suspension was incubated in an ultrasonic bath with sonication at 70°C for 15 minutes. One milliliter of the supernatant was then transferred to new tubes after centrifugation at 14,000 rpm (Eppendorf K-5418R). A total of 1.2 mL of a 10 mM sodium periodate solution (Merck, cat# 7790-28-5) was added to the suspension, and the mixture was shaken for 30 seconds. Then, 1.2 mL of a 0.2 M acetylacetone (Merck, cat# 123—54-6) solution was added to the former solution and kept in a water bath at 70°C for 10 minutes. The sample absorbance was measured with a UV – Vis – NIR spectrophotometer (Cary 5000, version 3.00, Agilent, Scan Version 6.2.0.1588), and the glycerol concentrations were estimated using the formula Y = 0.0055*Ab-0.0012. [Y = glycerol concentration; Ab = absorbance at 413 nm. The formula was build based on a twofold serious dilution of glycerol (>/= 99% purification, Merck, cat # 56-81-5). The glycerol density (microgram glycerol/mg hyphal weight) was calculated by dividing the total glycerol content of each sample by its corresponding hyphal weight.

#### Genomic DNA and RNA Extraction

The *C. militaris* conidia were inoculated in PDB media for 3 days at 130 rpm and 25°C. The hyphal pellets were washed with DEPC-treated water, and genomic DNA was extracted using the Monarch Genomic DNA Purification kit (New England Biolab, cat# T3010S). Total RNA was purified using the E.Z.N.A fungal RNA mini kit (Omega Bio- TeK, SKU R6840-01) – was used after TRIzol lysis (Thermo Fisher, cat # 15596026), and cDNA synthesis was performed using the ProtoScript® II First Strand cDNA Synthesis Kit (New England Biolab, cat# 6560S).

#### Phylogenetic Analysis

PCR amplification was performed using the primers ITS1 and ITS4 (**Additional File 5. Table S1**), and target DNA was amplified by OneTaq® 2X Master Mix with Standard Buffer (New England Biolab, cat# M0482S) under a thermal cycle of 94°C for 5 minutes, followed by 35 cycles of 94°C for 45 seconds, 52°C for 1 minute, 72°C for 1.5 minutes, and a final extension at 72°C for 5 minutes. The amplification products were purified with Qiaquick PCR Purification kits (Qiagen, cat # 28104) and then used directly for sequencing. Sequencing was performed using an ABI 3500 Series Genetic Analyzer (Thermo Fisher) and BigDye™ Terminator v3.1 Cycle Sequencing Kit (ThermoFisher, cat # 4337455). BioEdit software (43) was utilized to examine the DNA sequence quality and accuracy as well as select unambiguous bases. Multiple sequences were aligned, and BLAST search and phylogenetic analysis were performed using MEGA version 11.0.13 (44). The FASTA sequences of the *C. militaris* DM1066 (45) and JLCY-LI819 (46) strains were also included in the phylogenetic analysis with the minimum- evolutionary method (47) and interior-branch test (48). The neighbor joining tree (49) embedded on the NCBI website was used to explore the genetic relationships among *Ywt*, *Ydga* and other *C. militaris* strains.

#### Transcriptome Analysis

The transcriptome datasets from BioProject # PRJNA393201(50) with BioSample # YCCZ1-YCCZ6 (51)(22) were downloaded from the NCBI website using the SRA-tool kit (52). The fastq data were checked using FastQC (53), aligned to the *C. militaris* CM01 reference genome (54) using HISAT2 (55), and counted using HTSeq-count (56). Differential expression analysis was performed using EdgeR (57), and gene ontology enrichment analysis was performed using the ClusterProfiler package in R (58). All BioSamples, except for YCCZ3, had over 20 million reads with an overall alignment rate of at least 86%. YCCZ3 was excluded from further analysis. The expression patterns of the remaining samples were evaluated using hclust and plot functions in R. The plot showed that YCCZ5 (DG3) and YCCZ2 (DG1) clustered together and were farther from YCCZ1 (WT), while YCCZ4 (DG2) and YCCZ6 (DG4) were in the middle, between WT and the cluster of DG1 and DG3 (**Additional File 5. Figure S1**). DG1 is the second generation of WT and can still develop fruiting bodies. Therefore, we considered DG3 as the “true” *C. militaris* degenerate strain because it was the fifth passage of the WT and had lost its ability to form fruiting bodies (22). We compared it with the WT sample to identify differentially expressed genes.

#### RT–qPCR

To validate the dysregulated genes identified by Next Generation Sequencing and associated with phenotypic degeneration, we performed RT–qPCR using primers designed with PrimerQuest (59) from the idtdna.com website (**Additional File 5. Table S1**). Each RT–qPCR reaction had a total volume of 20 µl, containing cDNA, relevant primers, and the SYBR Green Real-time PCR Master Mix (Takara). Applied Biosystems™ 7500 Real-Time PCR Systems (Thermo Fisher) were used for RT–qPCR. Osh5 (CCM_00742 - oxysterol binding protein, putative) was used as an internal control (reference gene) for normalization of the target gene expression and to correct for variation between samples. The thermal cycle for RT-PCR was as follows: 94◦C for 1 min, followed by 40 cycles of 94°C for 45 seconds, 53°C for 60 seconds, and 72°C for 20 seconds. Melting curve analyses were performed at the end of each PCR reaction to ensure that only specific products were amplified. The comparative 2CT method (60) was employed to calculate relative expression levels among the target genes.

#### Statistical analysis

The mean ± standard error (SE) of replicates is presented for all data in this study. A one-tailed Student’s t test was conducted between the control and experimental groups to calculate the p values using the Student’s t test calculator provided by Socscistatistics.com. For comparison of radical expansion, spore density, fruiting body weight, and glycerol concentration, at least 5 biological replicates were used. For RT– qPCR evaluation, three technical replicates were used.

## Abbreviations

MAPK: Mitogen activated protein kinases
*RT–qPCR:*: Quantitative reverse transcription polymerase chain reaction
HOG: High-osmolarity glycerol
SF: Spore formation
PR: Pheromone Response
FG: Filamentous Growth
CWI: Cell wall integrity
ITS: Internal transcribed spacer
DG: Degenerate strain
WT: wild-type
NAC: N-acetyl cysteine
KCl: Potassium chloride
NaCl: Sodium Chloride
PDA: Potatoes dextrose agarose
CT: Control

## Acknowledgments

The authors thank Jennifer Curtis, William R. Folk and Shuqun Zhang for Manuscript review as well as Le Hai Yen and Vu Duy Nhan for the gift of the *C. militaris strains*.

## Availability of data and materials

All data generated or analyzed during this study are included in this published article (and its additional files). The RNAseq data associated with BioProject PRJNA393201 are deposited in NCBI databases by Yin at el., 2017(22). Accession numbers of this dataset can be found on the BioProject page at the following link: https://www.ncbi.nlm.nih.gov/bioproject/?term=PRJNA393201

## Author information

### Authors and Affiliations

National Institute for Technical Progress, C6 Thanh Xuan Bac, Thanh Xuan, Hanoi, Vietnam Chinh Quoc Hoang, Giang Huong Duong, Tao Xuan Vu, Tram Bao Tran.

Center for Biomedical Informatics, Vingroup Big Data Institute, and GeneStory JSC, Hanoi, Vietnam

Mai Hoang Tran

Department of Pharmacology and Biochemistry, National Institute of Medicinal Materials, and University of Medicine and Pharmacy, Vietnam National University, Ha Noi, Vietnam.

Hang Nguyet Thi Pham

### Contributions

CQH: conceptual design, data collection and analysis as well as manuscript preparation. GHTD and MHT: data collection and analysis. TXV, TBT, HNTP: conceptual design and data analysis. All authors read and approved the final version of the manuscript.

### Corresponding author

Correspondence to Chinh Q. Hoang.

## Ethics declarations

### Ethics approval and consent to participate

This section is not applicable.

### Consent for publication

This section is not applicable.

## Additional Files

### Additional File 1

**Table S1.**
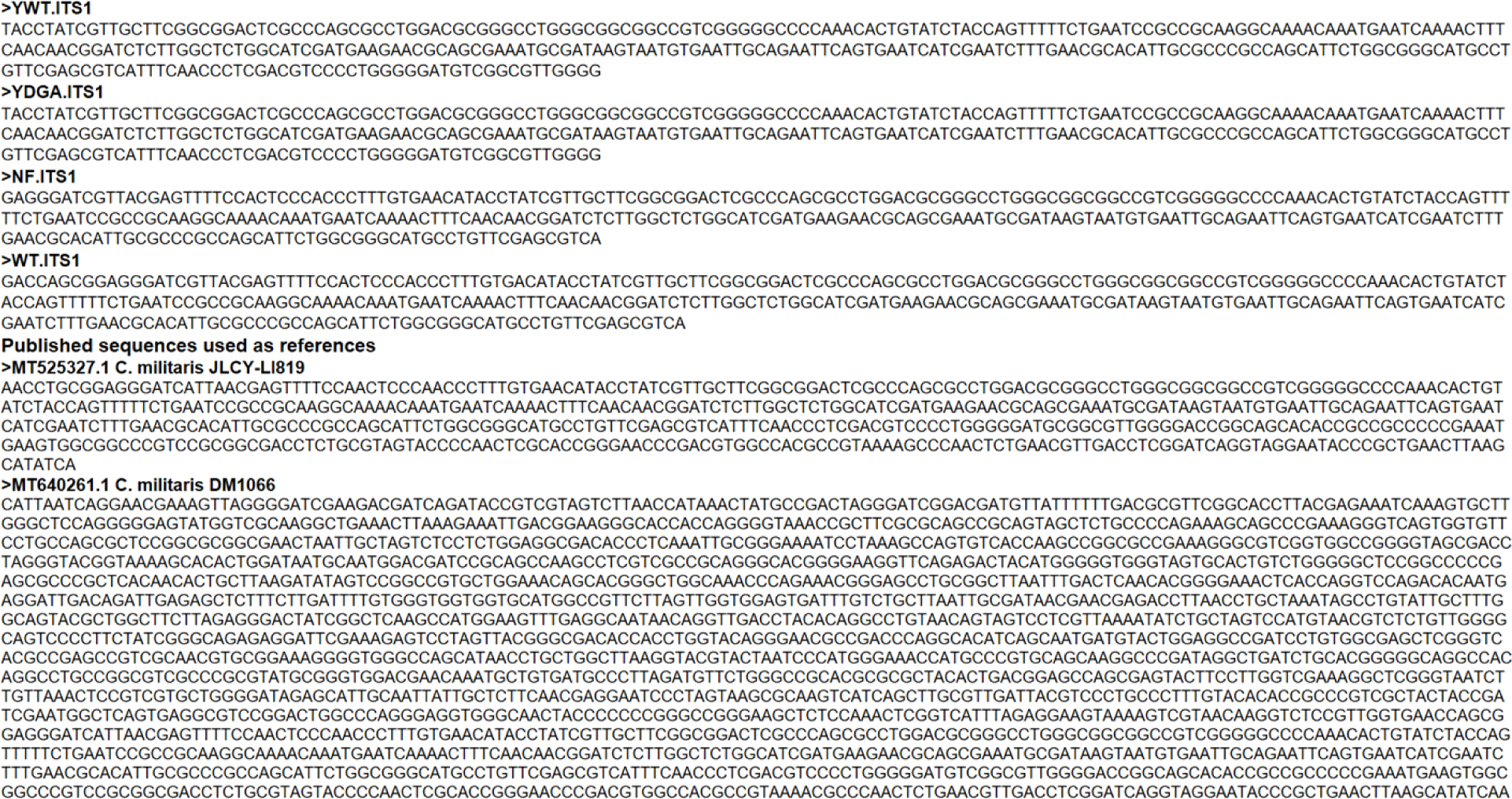
ITS sequences used for phylogenetic analysis.

**Figure S1.**
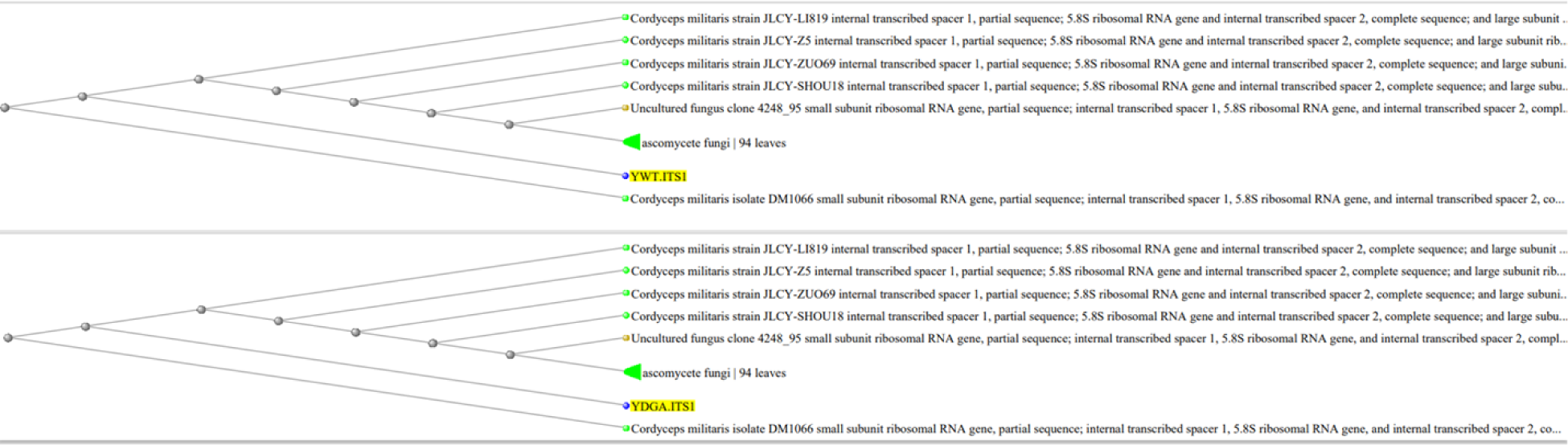
The neighbor joining tree analysis of the genetic relationship among *Ywt, Ydga*, and other *C. militaris* strains.

**Table S2.**
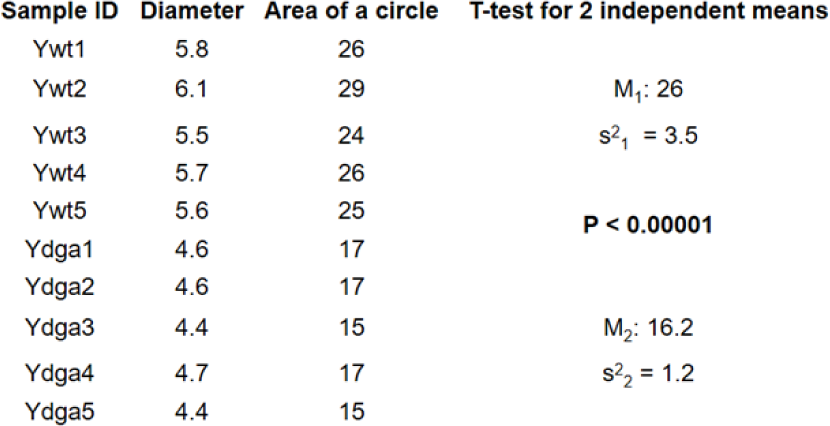
Radical expansion of C. militaris strain *Ywt* vs.Ydga.

| Sample ID | Diameter | Area of a circle | T-test for 2 independent means |
| --- | --- | --- | --- |
| Ywt1 | 5.8 | 26 | $M_1: 26$<br>$s^2_1 = 3.5$<br>**P < 0.00001** |
| Ywt2 | 6.1 | 29 |  |
| Ywt3 | 5.5 | 24 |  |
| Ywt4 | 5.7 | 26 |  |
| Ywt5 | 5.6 | 25 |  |
| Ydga1 | 4.6 | 17 | $M_2: 16.2$<br>$s^2_2 = 1.2$ |
| Ydga2 | 4.6 | 17 |  |
| Ydga3 | 4.4 | 15 |  |
| Ydga4 | 4.7 | 17 |  |
| Ydga5 | 4.4 | 15 |  |

**Table S3.** Spore density of *C. militairs* strain *Ywt vs. Ydga*.

| Sample ID | Count1 | Count2 | Count3 | Count4 | Count5 | Average | Dilution | Concentration | # ml | Total | Diameter | Area of circles | Density (10<sup>6</sup>/cm<sup>2</sup>) | T-test for 2 independent means |
| --- | --- | --- | --- | --- | --- | --- | --- | --- | --- | --- | --- | --- | --- | --- |
| Ywt1 | 46 | 45 | 53 | 45 | 43 | 46.4 | 1 | 11600000 | 8 | 92800000 | 6.2 | 30 | 3.07 | $M_1: 2.68$<br>$s^2_1 = 0.19$<br>**P < 0.00001** |
| Ywt2 | 42 | 43 | 49 | 56 | 38 | 45.6 | 1 | 11400000 | 8 | 91200000 | 6.4 | 32 | 2.83 |  |
| Ywt3 | 35 | 26 | 35 | 26 | 33 | 31 | 1 | 7750000 | 8 | 62000000 | 6.2 | 30 | 2.05 |  |
| Ywt4 | 33 | 40 | 40 | 32 | 43 | 37.6 | 1 | 9400000 | 8 | 75200000 | 6.3 | 31 | 2.41 |  |
| Ywt5 | 37 | 39 | 42 | 38 | 40 | 39.2 | 1 | 9800000 | 9 | 88200000 | 6.1 | 29 | 3.02 |  |
| Ydga1 | 136 | 224 | 160 | 152 | 156 | 165.6 | 1 | 41400000 | 7.6 | 314640000 | 4.7 | 17 | 18.14 | $M_2: 19.85$<br>$s^2_2 = 5.19$ |
| Ydga2 | 196 | 184 | 192 | 196 | 204 | 194.4 | 1 | 48600000 | 7.6 | 369360000 | 4.8 | 18 | 20.41 |  |
| Ydga3 | 184 | 176 | 160 | 260 | 252 | 206.4 | 1 | 51600000 | 7.6 | 392160000 | 4.6 | 17 | 23.60 |  |
| Ydga4 | 128 | 176 | 148 | 152 | 132 | 147.2 | 1 | 36800000 | 7.6 | 279680000 | 4.4 | 15 | 18.39 |  |
| Ydga5 | 130 | 160 | 168 | 150 | 140 | 149.6 | 1 | 37400000 | 7.6 | 284240000 | 4.4 | 15 | 18.69 |  |

### Additional File 2

**Table S1.** Biological pathways significantly enriched in downregulated genes.

| Pathway ID | Pathway names | GeneRatio | BgRatio | pvalue | p.adjust | qvalue | # of DE genes |
| --- | --- | --- | --- | --- | --- | --- | --- |
| cmt02010 | ABC transporters - Cordyceps militaris | 16/152 | 38/2057 | 1.97E-09 | 1.38E-07 | 1.35E-07 | 16 |
| cmt04011 | MAPK signaling pathway - yeast - Cordyceps militaris | 16/152 | 63/2057 | 6.36E-06 | 0.000223 | 0.0002177 | 16 |
| cmt00520 | Amino sugar and nucleotide sugar metabolism - Cordyceps militaris | 13/152 | 57/2057 | 0.000166 | 0.003872 | 0.00378511 | 13 |

| No. | DE genes in ABC transporter | DE genes in MAPK pathway | DE genes in Amino sugar and nucleotide sugar metabolism |
| --- | --- | --- | --- |
| 1 | CCM_08739 | CCM_05444 | CCM_06535 |
| 2 | CCM_03607 | CCM_03130 | CCM_08231 |
| 3 | CCM_01696 | CCM_06568 | CCM_02966 |
| 4 | CCM_02004 | CCM_02415 | CCM_00318 |
| 5 | CCM_02386 | CCM_07679 | CCM_02965 |
| 6 | CCM_06618 | CCM_07878 | CCM_07033 |
| 7 | CCM_01467 | CCM_08862 | CCM_05430 |
| 8 | CCM_00623 | CCM_02506 | CCM_08511 |
| 9 | CCM_03253 | CCM_09124 | CCM_04624 |
| 10 | CCM_04460 | CCM_01444 | CCM_01980 |
| 11 | CCM_06149 | CCM_07079 | CCM_05688 |
| 12 | CCM_03445 | CCM_01361 | CCM_04625 |
| 13 | CCM_07363 | CCM_08816 | CCM_06973 |
| 14 | CCM_08649 | CCM_02193 |  |
| 15 | CCM_06538 | CCM_05107 |  |
| 16 | CCM_08294 | CCM_09157 |  |

**Table S2.** The *C. militaris Ydga* strain has higher intracellular glycerol contents than that of *Ywt* strain.

| Sample ID | OD | Glycerol content | dilution | Total glycerol content | *C. militaris* wet weight | Glycerol content/g of *C.militaris* | T-test for 2 independent means |
| --- | --- | --- | --- | --- | --- | --- | --- |
| ywt1 | 0.846 | 0.003453 | 1 | 0.003453 | 7.1 | 0.486338028 | $M_1: 1.09$<br>$s^2_1 = 0.13$ |
| ywt2 | 3.102 | 0.015861 | 1 | 0.015861 | 10.9 | 1.455137615 |  |
| ywt3 | 2.621 | 0.0132155 | 1 | 0.0132155 | 12 | 1.101291667 |  |
| ywt4 | 3.484 | 0.017962 | 1 | 0.017962 | 15 | 1.197466667 |  |
| ywt5 | 4.017 | 0.0208935 | 1 | 0.0208935 | 17.2 | 1.214738372 |  |
| | | | | | | | $p < 0.001$ |
| ydga1 | 4.064 | 0.021152 | 1 | 0.021152 | 10.2 | 2.07372549 | $M_2: 1.93$<br>$s^2_2 = 0.01$ |
| ydga2 | 4 | 0.0208 | 1 | 0.0208 | 10.4 | 2 |  |
| ydga3 | 4.337 | 0.0226535 | 1 | 0.0226535 | 11.6 | 1.952887931 |  |
| ydga4 | 4.023 | 0.0209265 | 1 | 0.0209265 | 11.9 | 1.758529412 |  |
| ydga5 | 4.521 | 0.0236655 | 1 | 0.0236655 | 12.7 | 1.863425197 |  |

OD = Optical density at 413 nm

### Additional File 3

**Table S1.** the CWI stressor represses the radical expansion and sporulation of C. militaris strain *Ywt vs. Ydga*.

| Sample ID | Areas of Circles | T-test for 2 independent means | Spore density (10<sup>6</sup>) | T-test for 2 independent means | Sample ID | Areas of Circles | T-test for 2 independent means | Spore density (10<sup>6</sup>) | T-test for 2 independent means |
| --- | --- | --- | --- | --- | --- | --- | --- | --- | --- |
| Ywt.PDA1 | 22.90 |  | 1.21 |  | Ydga.PDA1 | 15.21 |  | 44.33 |  |
| Ywt.PDA2 | 21.24 | M<sub>1</sub>: 23.26 | 1.01 | M<sub>1</sub>: 1.03 | Ydga.PDA2 | 15.21 | M<sub>1</sub>: 15.35 | 44.72 | M<sub>1</sub>: 39.05 |
| Ywt.PDA3 | 24.63 | s<sup>2</sup><sub>1</sub> = 1.65 | 0.91 | s<sup>2</sup><sub>1</sub> = 0.01 | Ydga.PDA3 | 15.90 | s<sup>2</sup><sub>1</sub> = 0.1 | 33.20 | s<sup>2</sup><sub>1</sub> = 28.16 |
| Ywt.PDA4 | 23.76 |  | 0.98 |  | Ydga.PDA4 | 15.21 |  | 34.86 |  |
| Ywt.PDA5 | 23.76 |  | 1.05 |  | Ydga.PDA5 | 15.21 |  | 38.14 |  |
| **PDA vs. CR** |  | **p < 0.0001** |  | **p < 0.001** |  |  | **P < 0.0001** |  | **P = 0.006** |
| Ywt.CR1 | 13.20 |  | 0.61 |  | Ydga.CR1 | 11.95 |  | 26.62 |  |
| Ywt.CR2 | 13.20 |  | 0.74 |  | Ydga.CR2 | 11.34 |  | 27.51 |  |
| Ywt.CR3 | 15.21 | M<sub>2</sub>: 14.4 | 0.64 | M<sub>2</sub>: 0.65 | Ydga.CR3 | 11.34 | M<sub>2</sub>: 11.23 | 31.21 | M<sub>2</sub>: 30.35 |
| Ywt.CR4 | 15.21 | s<sup>2</sup><sub>2</sub> = 1.2 | 0.70 | s<sup>2</sup><sub>2</sub> = 0.01 | Ydga.CR4 | 10.75 | s<sup>2</sup><sub>2</sub> = 0.25 | 33.48 | s<sup>2</sup><sub>2</sub> = 9.79 |
| Ywt.CR5 | 15.21 |  | 0.53 |  | Ydga.CR5 | 10.75 |  | 32.92 |  |

PDA = Potatoes Dextrose Agar; CR = PDA + Congo Red (200 ug/ml)

**Table S2.** The oxidative stressor slightly suppresses the radical expansion, but promotes sporulation of *C. militaris* strain *Ywt vs. Ydga*.

| Sample ID | Areas of Circles | T-test for 2 independent means | Spore density (10<sup>6</sup>) | T-test for 2 independent means | Sample ID | Areas of Circles | T-test for 2 independent means | Spore density (10<sup>6</sup>) | T-test for 2 independent means |
| --- | --- | --- | --- | --- | --- | --- | --- | --- | --- |
| Ywt.PDA1 | 26.42 |  | 0.98 |  | Ydga. PDA1 | 15.90 |  | 45.27 |  |
| Ywt.PDA2 | 25.52 | M<sub>1</sub>: 26.61 | 1.46 | M<sub>1</sub>: 1.3 | Ydga. PDA2 | 16.62 | M<sub>1</sub>: 16.48 | 62.87 | M<sub>1</sub>: 60.49 |
| Ywt.PDA3 | 28.27 | s<sup>2</sup><sub>1</sub> = 1.02 | 1.41 | s<sup>2</sup><sub>1</sub> = 0.04 | Ydga. PDA3 | 17.35 | s<sup>2</sup><sub>1</sub> = 0.36 | 69.08 | s<sup>2</sup><sub>1</sub> = 92.45 |
| Ywt.PDA4 | 26.42 |  | 1.43 |  | Ydga. PDA4 | 15.90 |  | 57.64 |  |
| Ywt.PDA5 | 26.42 |  | 1.21 |  | Ydga. PDA5 | 16.62 |  | 67.60 |  |
| **PDA vs. H2O2** |  | **p = 0.003** |  | **p < 0.001** |  |  | **p < 0.05** |  | **p < 0.05** |
| Ywt.H2O2.1 | 21.24 |  | 1.54 |  | Ydga.H2O2.1 | 13.20 |  | 57.26 |  |
| Ywt.H2O2.2 | 24.63 |  | 1.94 |  | Ydga.H2O2.2 | 13.20 |  | 71.14 |  |
| Ywt.H2O2.3 | 24.63 | M<sub>2</sub>: 23.27 | 1.94 | M<sub>2</sub>: 1.96 | Ydga.H2O2.3 | 13.85 | M<sub>2</sub>: 13.46 | 86.40 | M<sub>2</sub>: 73.08 |
| Ywt.H2O2.4 | 21.24 | s<sup>2</sup><sub>2</sub> = 3.45 | 2.30 | s<sup>2</sup><sub>2</sub> = 0.08 | Ydga.H2O2.4 | 13.85 | s<sup>2</sup><sub>2</sub> = 0.13 | 77.30 | s<sup>2</sup><sub>2</sub> = 112.31 |
| Ywt.H2O2.5 | 24.63 |  | 2.10 |  | Ydga.H2O2.5 | 13.20 |  | 73.28 |  |
| **PDA vs. NAC** |  | **p = 0.0002** |  | **p > 0.05** |  |  | **p = 0.005** |  | **p > 0.05** |
| Ywt.NAC1 | 23.76 |  | 0.98 |  | Ydga.NAC1 | 15.21 |  | 32.49 |  |
| Ywt.NAC2 | 23.76 |  | 1.06 |  | Ydga.NAC2 | 15.21 |  | 44.98 |  |
| Ywt.NAC3 | 24.63 | M<sub>2</sub>: 23.93 | 1.27 | M<sub>2</sub>: 1.11 | Ydga.NAC3 | 13.85 | M<sub>2</sub>: 15.07 | 45.12 | M<sub>2</sub>: 51.2 |
| Ywt.NAC4 | 23.76 | s<sup>2</sup><sub>2</sub> = 0.15 | 1.06 | s<sup>2</sup><sub>2</sub> = 0.06 | Ydga.NAC4 | 15.90 | s<sup>2</sup><sub>2</sub> = 0.56 | 67.20 | s<sup>2</sup><sub>2</sub> = 226.87 |
| Ywt.NAC5 | 23.76 |  | 1.20 |  | Ydga.NAC5 | 15.21 |  | 66.23 |  |

PDA = Potatoes Dextrose Agar; H2O2 = PDA + H2O2 (0.04%); NAC = PDA + N-AcetylCysteine (200 mM)

**Table S3.** Hyperosmotic stressors suppress the radical expansion, but promotes sporulation of *C. militaris* strain *Ywt vs. Ydga*.

| Sample ID | Areas of Circles | T-test for 2 independent means | Spore density (10<sup>6</sup>) | T-test for 2 independent means | Sample ID | Areas of Circles | T-test for 2 independent means | Spore density (10<sup>6</sup>) | T-test for 2 independent means |
| --- | --- | --- | --- | --- | --- | --- | --- | --- | --- |
| Ywt.PDA1 | 22.06183 |  | 0.781893 |  | Ydga.PDA1 | 15.20531 |  | 25.35299 |  |
| Ywt.PDA2 | 24.63009 | M<sub>1</sub>: 23.42 | 0.669912 | M<sub>1</sub>: 0.68 | Ydga.PDA2 | 16.61903 | M<sub>1</sub>: 16.63 | 20.75934 | M<sub>1</sub>: 24.79 |
| Ywt.PDA3 | 22.90221 | s<sub>1</sub><sup>2</sup> = 0.95 | 0.534883 | s<sub>1</sub><sup>2</sup> = 0.01 | Ydga.PDA3 | 15.90431 | s<sub>1</sub><sup>2</sup> = 1.31 | 29.14304 | s<sub>1</sub><sup>2</sup> = 8.99 |
| Ywt.PDA4 | 23.75829 |  | 0.694494 |  | Ydga.PDA4 | 18.09557 |  | 24.17718 |  |
| Ywt.PDA5 | 23.75829 |  | 0.736585 |  | Ydga.PDA5 | 17.34945 |  | 24.49646 |  |
| **PDA vs. NaCl** |  | **p < 0.00001** |  | **p < 0.00001** |  |  | **p < 0.00001** |  | **p < 0.00001** |
| Ywt.NaCl1 | 9.621128 |  | 4.859098 |  | Ydga.NaCl1 | 3.141593 |  | 44.08592 |  |
| Ywt.NaCl2 | 10.17876 |  | 5.010433 |  | Ydga.NaCl2 | 3.141593 |  | 49.17888 |  |
| Ywt.NaCl3 | 9.621128 |  | 5.794539 |  | Ydga.NaCl3 | 5.725553 |  | 44.97382 |  |
| Ywt.NaCl4 | 8.552986 | M<sub>2</sub>: 9.41 | 5.027484 | M<sub>2</sub>: 5.12 | Ydga.NaCl4 | 4.908739 | M<sub>2</sub>: 4.08 | 50.11471 | M<sub>2</sub>: 47.26 |
| Ywt.NaCl5 | 9.079203 | s<sub>2</sub><sup>2</sup> = 0.38 | 4.928847 | s<sub>2</sub><sup>2</sup> = 0.15 | Ydga.NaCl5 | 3.463606 | s<sub>2</sub><sup>2</sup> = 1.38 | 47.92693 | s<sub>2</sub><sup>2</sup> = 6.89 |
| **PDA vs. KCl** |  | **p < 0.00001** |  | **p < 0.00001** |  |  | **p < 0.00001** |  | **p = 0.002** |
| Ywt.KCl1 | 8.042477 |  | 5.632593 |  | Ydga.KCl1 | 5.309292 |  | 45.76882 |  |
| Ywt.KCl2 | 10.17876 |  | 5.467267 |  | Ydga.KCl2 | 5.309292 |  | 41.81349 |  |
| Ywt.KCl3 | 9.621128 |  | 6.319426 |  | Ydga.KCl3 | 6.157522 |  | 35.72866 |  |
| Ywt.KCl4 | 9.079203 | M<sub>2</sub>: 9.09 | 6.542425 | M<sub>2</sub>: 6.04 | Ydga.KCl4 | 5.725553 | M<sub>2</sub>: 5.73 | 31.52534 | M<sub>2</sub>: 37.22 |
| Ywt.KCl5 | 8.552986 | s<sub>1</sub><sup>2</sup> = 0.71 | 6.260971 | s<sub>1</sub><sup>2</sup> = 0.22 | Ydga.KCl5 | 6.157522 | s<sub>1</sub><sup>2</sup> = 0.18 | 31.26258 | s<sub>1</sub><sup>2</sup> = 41.08 |

PDA = Potatoes Dextrose Agar; NaCl = PDA + NaCl(0.4 M); KCl = PDA + KCl(0.4 M)

**Table S4.** Hyperosmotic stressors increase the intracellular glycerol contents of *C. militaris* strain *Ywt*.

| Sample ID | OD | Glycerol content | dilution | Total glycerol content | *C. militaris* wet weight | Glycerol content/g of *C.militaris* | T-test for 2 independent means |
| --- | --- | --- | --- | --- | --- | --- | --- |
| Ywt.PDA1 | 2.4559 | 0.0123 | 1.0000 | 0.0123 | 8.5000 | 1.4480 |  |
| Ywt.PDA2 | 2.8176 | 0.0143 | 1.0000 | 0.0143 | 9.1000 | 1.5711 | M<sub>1</sub>: 1.66 |
| Ywt.PDA3 | 3.3583 | 0.0173 | 1.0000 | 0.0173 | 9.1000 | 1.8979 | s<sub>1</sub><sup>2</sup> = 0.06 |
| Ywt.PDA4 | 3.9728 | 0.0207 | 1.0000 | 0.0207 | 14.4000 | 1.4341 |  |
| Ywt.PDA5 | 2.7768 | 0.0141 | 2.0000 | 0.0281 | 14.4000 | 1.9545 |  |
| **PDA vs. Sor** |  |  |  |  |  |  | **p < 0.00001** |
| Ywt.Sor1 | 3.6380 | 0.0188 | 1.0000 | 0.0188 | 5.2000 | 3.6171 |  |
| Ywt.Sor2 | 3.0462 | 0.0156 | 2.0000 | 0.0311 | 6.7000 | 4.6431 | M<sub>1</sub>: 4.45 |
| Ywt.Sor3 | 3.1122 | 0.0159 | 2.0000 | 0.0318 | 6.9000 | 4.6137 | s<sub>1</sub><sup>2</sup> = 0.22 |
| Ywt.Sor4 | 3.3454 | 0.0172 | 2.0000 | 0.0344 | 7.2000 | 4.7777 |  |
| Ywt.Sor5 | 3.2122 | 0.0165 | 2.0000 | 0.0329 | 7.2000 | 4.5743 |  |
| **PDA vs. NaCl** |  |  |  |  |  |  | **p < 0.00001** |
| Ywt.NaCl1 | 2.6771 | 0.0135 | 3.0000 | 0.0406 | 8.1000 | 5.0089 |  |
| Ywt.NaCl2 | 2.3887 | 0.0119 | 4.0000 | 0.0478 | 8.1000 | 5.8952 | M<sub>1</sub>: 5.47 |
| Ywt.NaCl3 | 3.2361 | 0.0166 | 3.0000 | 0.0498 | 8.2000 | 6.0726 | s<sub>1</sub><sup>2</sup> = 0.64 |
| Ywt.NaCl4 | 3.2837 | 0.0169 | 3.0000 | 0.0506 | 8.3000 | 6.0941 |  |
| Ywt.NaCl5 | 2.9473 | 0.0150 | 3.0000 | 0.0450 | 10.5000 | 4.2886 |  |
| **PDA vs. KCl** |  |  |  |  |  |  | **p < 0.00001** |
| Ywt.KCl1 | 2.4435 | 0.0122 | 3.0000 | 0.0367 | 6.5000 | 5.6489 |  |
| Ywt.KCl2 | 3.2313 | 0.0166 | 3.0000 | 0.0497 | 8.1000 | 6.1378 | M<sub>1</sub>: 5.29 |
| Ywt.KCl3 | 3.4893 | 0.0180 | 3.0000 | 0.0540 | 10.4000 | 5.1897 | s<sub>1</sub><sup>2</sup> = 0.42 |
| Ywt.KCl4 | 3.3073 | 0.0170 | 3.0000 | 0.0510 | 11.5000 | 4.4321 |  |
| Ywt.KCl5 | 3.1073 | 0.0159 | 3.0000 | 0.0477 | 9.5000 | 5.0179 |  |

PDA = Potatoes Dextrose Agar; Sor = PDA + Sorbitol (1 M), NaCl = PDA + NaCl(0.4 M); KCl = PDA + KCl(0.4 M), OD = Optical density at 413 nm

**Table S5.** Hyperosmotic stressors increase intracellular glycerol contents of *C. militaris* strain Ydga.

| Sample ID | OD | Glycerol content | dilution | Total glycerol content | *C. militaris* wet weight | Glycerol content/g of *C. militaris* | T-test for 2 independent means |
| --- | --- | --- | --- | --- | --- | --- | --- |
| Ydga.PDA1 | 3.1610 | 0.0162 | 2.0000 | 0.0324 | 8.1000 | 3.9964 |  |
| Ydga.PDA2 | 2.8896 | 0.0147 | 2.0000 | 0.0294 | 8.3000 | 3.5404 | $M_1: 3.17$ |
| Ydga.PDA3 | 2.8404 | 0.0144 | 2.0000 | 0.0288 | 8.4000 | 3.4338 | $s^2_1 = 0.5$ |
| Ydga.PDA4 | 2.7218 | 0.0138 | 2.0000 | 0.0275 | 11.4000 | 2.4158 |  |
| Ydga.PDA5 | 2.8208 | 0.0143 | 2.0000 | 0.0286 | 11.7000 | 2.4470 |  |
| **PDA vs. Sor** |  |  |  |  |  |  | **p < 0.0001** |
| Ydga.Sor1 | 3.0619 | 0.0156 | 2.0000 | 0.0313 | 4.0000 | 7.8203 |  |
| Ydga.Sor2 | 3.7059 | 0.0192 | 2.0000 | 0.0384 | 4.4000 | 8.7193 |  |
| Ydga.Sor3 | 3.5516 | 0.0183 | 2.0000 | 0.0367 | 5.5000 | 6.6668 | $M_2: 7.11$ |
| Ydga.Sor4 | 3.0516 | 0.0156 | 2.0000 | 0.0312 | 5.0000 | 6.2334 | $s^2_2 = 1.26$ |
| Ydga.Sor5 | 3.5516 | 0.0183 | 2.0000 | 0.0367 | 6.0000 | 6.1112 |  |
| **PDA vs. NaCl** |  |  |  |  |  |  | **p < 0.0001** |
| Ydga.NaCl1 | 2.7985 | 0.0142 | 4.0000 | 0.0568 | 5.3000 | 10.7108 |  |
| Ydga.NaCl2 | 1.7036 | 0.0082 | 4.0000 | 0.0327 | 5.4000 | 6.0515 | $M_2: 9.01$ |
| Ydga.NaCl3 | 2.5367 | 0.0128 | 4.0000 | 0.0510 | 5.7000 | 8.9489 | $s^2_2 = 3.14$ |
| Ydga.NaCl4 | 2.4367 | 0.0122 | 4.0000 | 0.0488 | 5.0000 | 9.7617 |  |
| Ydga.NaCl5 | 2.8367 | 0.0144 | 4.0000 | 0.0576 | 6.0000 | 9.6014 |  |
| **PDA vs. KCl** |  |  |  |  |  |  | **p < 0.00001** |
| Ydga.KCl1 | 3.9778 | 0.0207 | 4.0000 | 0.0827 | 6.5000 | 12.7248 |  |
| Ydga.KCl2 | 4.2072 | 0.0219 | 4.0000 | 0.0878 | 6.2000 | 14.1547 | $M_2: 12.23$ |
| Ydga.KCl3 | 4.0186 | 0.0209 | 4.0000 | 0.0836 | 6.1000 | 13.7064 | $s^2_2 = 3.47$ |
| Ydga.KCl4 | 2.4433 | 0.0122 | 4.0000 | 0.0490 | 4.6000 | 10.6418 |  |
| Ydga.KCl5 | 3.2014 | 0.0164 | 4.0000 | 0.0656 | 6.6000 | 9.9442 |  |

PDA = Potatoes Dextrose Agar; Sor = PDA + Sorbitol (1 M); NaCl = PDA + NaCl(0.4 M); KCl = PDA + KCl(0.4 M); OD = Optical density at 413 nm

### Additional File 4

**Table S1.** Hyperosmotic stress suppresses the radical expansion, but increases sporulation of C. militaris strain *Wt, Nt, Ywt* and *Ydga*.

| Sample ID | Areas of Circles | T-test for 2 independent means | Spore density (10<sup>6</sup>) | T-test for 2 independent means | Sample ID | Areas of Circles | T-test for 2 independent means | Spore density (10<sup>6</sup>) | T-test for 2 independent means |
| --- | --- | --- | --- | --- | --- | --- | --- | --- | --- |
| Wt.PDA1 | 37.26 |  | 3.64 |  | Nf.PDA1 | 39.79 |  | 1.05 |  |
| Wt.PDA2 | 41.08 | M<sub>1</sub>: 39.54 | 2.54 | M<sub>1</sub>: 3.42 | Nf.PDA2 | 42.39 | M<sub>1</sub>: 39.32 | 2.18 | M<sub>1</sub>: 1.34 |
| Wt.PDA3 | 41.08 | s<sub>1</sub><sup>2</sup> = 2.77 | 4.73 | s<sub>1</sub><sup>2</sup> = 0.084 | Nf.PDA3 | 39.79 | s<sub>1</sub><sup>2</sup> = 7.58 | 1.15 | s<sub>1</sub><sup>2</sup> = 0.22 |
| Wt.PDA4 | 39.79 |  | 2.54 |  | Nf.PDA4 | 34.82 |  | 1.18 |  |
| Wt.PDA5 | 38.51 |  | 3.64 |  | Nf.PDA5 | 39.79 |  | 1.15 |  |
| Wt.Kcl1 | 20.04 | p < 0.00001 | 24.09 | p < 0.00001 | Nf.Kcl1 | 17.40 | P < 0.00001 | 12.36 | P < 0.0001 |
| Wt.Kcl2 | 19.14 |  | 24.23 |  | Nf.Kcl2 | 18.26 |  | 23.78 |  |
| Wt.Kcl3 | 20.04 | M<sub>2</sub>: 20.04 | 20.85 | M<sub>2</sub>: 22.5 | Nf.Kcl3 | 17.40 | M<sub>2</sub>: 17.92 | 18.44 | M<sub>2</sub>: 17.86 |
| Wt.Kcl4 | 20.96 | s<sub>2</sub><sup>2</sup> = 0.41 | 22.09 | s<sub>2</sub><sup>2</sup> = 2.51 | Nf.Kcl4 | 18.26 | s<sub>2</sub><sup>2</sup> = 0.22 | 15.15 | s<sub>2</sub><sup>2</sup> = 18.97 |
| Wt.Kcl5 | 20.04 |  | 21.23 |  | Nf.Kcl5 | 18.26 |  | 19.56 |  |
| Ywt.PDA1 | 39.79 |  | 2.33 |  | Ydga.PDA1 | 22.86 |  | 13.76 |  |
| Ywt.PDA2 | 42.39 | M<sub>1</sub>: 40.57 | 2.15 | M<sub>1</sub>: 1.97 | Ydga.PDA2 | 23.85 | M<sub>1</sub>: 22.11 | 15.49 | M<sub>1</sub>: 2.8 |
| Ywt.PDA3 | 39.79 | s<sub>1</sub><sup>2</sup> = 1.35 | 1.56 | s<sub>1</sub><sup>2</sup> = 0.09 | Ydga.PDA3 | 21.90 | s<sub>1</sub><sup>2</sup> = 1.99 | 17.91 | s<sub>1</sub><sup>2</sup> = 28.16 |
| Ywt.PDA4 | 41.08 |  | 1.83 |  | Ydga.PDA4 | 20.04 |  | 13.96 |  |
| Ywt.PDA5 | 39.79 |  | 1.98 |  | Ydga.PDA5 | 21.90 |  | 14.80 |  |
| Ywt.Kcl1 | 16.56 | p < 0.00001 | 21.98 | p < 0.00001 | Ydga.Kcl1 | 13.41 | P < 0.00001 | 26.06 | P = 0.0003 |
| Ywt.Kcl2 | 15.74 |  | 18.72 |  | Ydga.Kcl2 | 14.95 |  | 38.11 |  |
| Ywt.Kcl3 | 17.40 | M<sub>2</sub>: 16.73 | 15.13 | M<sub>2</sub>: 18.53 | Ydga.Kcl3 | 13.41 | M<sub>2</sub>: 13.57 | 23.57 | M<sub>2</sub>: 29.51 |
| Ywt.Kcl4 | 17.40 | s<sub>2</sub><sup>2</sup> = 0.48 | 17.75 | s<sub>2</sub><sup>2</sup> = 6.1 | Ydga.Kcl4 | 12.68 | s<sub>2</sub><sup>2</sup> = 0.69 | 31.53 | s<sub>2</sub><sup>2</sup> = 31.67 |
| Ywt.Kcl5 | 16.56 |  | 19.08 |  | Ydga.Kcl5 | 13.41 |  | 28.30 |  |

PDA = Potatoes Dextrose Agar; Kcl = PDA + Kcl (0.4 M)

**Table S2.** Hyperosmotic stress increases the intracellular glycerol contents of C. militaris strain *Wt, Nt, Ywt* and *Ydga*.

| Sample ID | OD | Glycerol contents | Glycerol content/g of *C.militaris* | T-test for 2 independent means | Sample ID | OD | Glycerol contents | Glycerol content/g of *C.militaris* | T-test for 2 independent means |
| --- | --- | --- | --- | --- | --- | --- | --- | --- | --- |
| Wt.PDA1 | 0.7449 | 0.002897 | 0.44232 |  | Nf.PDA1 | 2.8868 | 0.0147 | 0.9439 |  |
| Wt.PDA2 | 0.8642 | 0.003553 | 0.51125 | M<sub>1</sub>: 0.46 | Nf.PDA2 | 2.863 | 0.0145 | 0.7982 | M<sub>1</sub>: 0.83 |
| Wt.PDA3 | 0.6072 | 0.00214 | 0.40755 | s<sub>1</sub><sup>2</sup> = 0 | Nf.PDA3 | 3.8645 | 0.0201 | 0.8062 | s<sub>1</sub><sup>2</sup> = 0.02 |
| Wt.PDA4 | 0.7072 | 0.00269 | 0.43035 |  | Nf.PDA4 | 3.02 | 0.0154 | 0.7783 |  |
| Wt.PDA5 | 0.9072 | 0.00379 | 0.52271 |  | Nf.PDA5 | 3.21 | 0.0165 | 0.7998 |  |
| Wt.Kcl1 | 4.0125 | 0.020869 | 7.19613 | p < 0.00001 | Nf.Kcl1 | 3.1202 | 0.016 | 3.1373 | P < 0.0001 |
| Wt.Kcl2 | 4.1472 | 0.021609 | 8.35149 |  | Nf.Kcl2 | 3.2697 | 0.0168 | 3.1444 |  |
| Wt.Kcl3 | 3.8292 | 0.019861 | 6.11099 | M<sub>2</sub>: 7.45 | Nf.Kcl3 | 3.2883 | 0.0169 | 2.4832 | M<sub>2</sub>: 2.94 |
| Wt.Kcl4 | 3.6149 | 0.018682 | 7.66448 | s<sub>2</sub><sup>2</sup> = 0.73 | Nf.Kcl4 | 3.3797 | 0.0174 | 3.0373 | s<sub>2</sub><sup>2</sup> = 0.07 |
| Wt.Kcl5 | 3.8149 | 0.019782 | 7.91287 |  | Nf.Kcl5 | 3.4883 | 0.018 | 2.9067 |  |
| Ywt.PDA1 | 0.3297 | 0.000613 | 0.07031 |  | Ydga.PDA1 | 1.3 | 0.0059 | 0.1245 |  |
| Ywt.PDA2 | 0.3594 | 0.000777 | 0.07966 | M<sub>1</sub>: 0.08 | Ydga.PDA2 | 1.5595 | 0.0074 | 0.5589 | M<sub>1</sub>: 0.62 |
| Ywt.PDA3 | 0.3865 | 0.000926 | 0.08859 | s<sub>1</sub><sup>2</sup> = 0 | Ydga.PDA3 | 2.309 | 0.0115 | 0.7628 | s<sub>1</sub><sup>2</sup> = 0.09 |
| Ywt.PDA4 | 0.3665 | 0.000816 | 0.08819 |  | Ydga.PDA4 | 2.159 | 0.0107 | 0.8455 |  |
| Ywt.PDA5 | 0.3565 | 0.000761 | 0.08694 |  | Ydga.PDA5 | 2.209 | 0.0109 | 0.7877 |  |
| Ywt.Kcl1 | 3.1137 | 0.015925 | 3.30918 | p < 0.00001 | Ydga.Kcl1 | 2.7912 | 0.0142 | 3.4621 | p < 0.00001 |
| Ywt.Kcl2 | 3.1856 | 0.016321 | 3.05062 |  | Ydga.Kcl2 | 2.6105 | 0.0132 | 3.0161 |  |
| Ywt.Kcl3 | 4.9872 | 0.02623 | 3.04114 | M<sub>2</sub>: 3.2 | Ydga.Kcl3 | 2.3643 | 0.0118 | 2.5591 | M<sub>2</sub>: 2.99 |
| Ywt.Kcl4 | 4.3872 | 0.02293 | 3.47421 | s<sub>2</sub><sup>2</sup> = 0.14 | Ydga.Kcl4 | 2.4643 | 0.0124 | 2.959 | s<sub>2</sub><sup>2</sup> = 0.1 |
| Ywt.Kcl5 | 4.6872 | 0.02458 | 3.14621 |  | Ydga.Kcl5 | 2.6243 | 0.0132 | 2.9573 |  |

PDA = Potatoes Dextrose Agar; Kcl = PDA + Kcl (0.4 M); OD = Optical density at 413 nm

**Table S3.** Hyperosmotic stress represses the fruiting body development of *C. militaris* strain *Wt, Nt, Ywt*.

| Sample ID | FB dry weights | T-test for 2 independent means | Sample ID | FB dry weights | T-test for 2 independent means | Sample ID | FB dry weights | T-test for 2 independent means |
| --- | --- | --- | --- | --- | --- | --- | --- | --- |
| Ywt.DC1 | 3.08 |  | Wt.DC1 | 2.44 |  | Nf.DC1 | 2.38 |  |
| Ywt.DC2 | 3.15 | | Wt.DC2 | 2.67 | | Nf.DC2 | 2.53 | $M_1: 3.2$ |
| Ywt.DC3 | 3.47 | $M_1: 3.58$ | Wt.DC3 | 2.53 | $M_1: 2.73$ | Nf.DC3 | 3.23 | $s^2_1 = 0.33$ |
| Ywt.DC4 | 3.62 | $s^2_1 = 0.12$ | Wt.DC4 | 3.03 | $s^2_1 = 0.27$ | Nf.DC4 | 3.31 | |
| Ywt.DC5 | 3.69 |  | Wt.DC5 | 2.21 |  | Nf.DC5 | 3.36 |  |
| Ywt.DC6 | 3.71 |  | Wt.DC6 | 2.48 |  | Nf.DC6 | 4.02 |  |
| Ywt.DC7 | 3.73 | | Wt.DC7 | 3.75 | | Nf.DC7 | 3.58 | $p = 0.0007$ |
| Ywt.DC8 | 4.16 | $p < 0.00001$ | Wt.Kcl1 | 1.58 | $p = 0.002$ | Nf.Kcl1 | 1.93 | |
| Ywt.Kcl1 | 1.5 |  | Wt.Kcl2 | 2.16 |  | Nf.Kcl2 | 2.06 |  |
| Ywt.Kcl2 | 1.55 |  | Wt.Kcl3 | 0.71 |  | Nf.Kcl3 | 2.27 |  |
| Ywt.Kcl3 | 1.68 | | Wt.Kcl4 | 0.48 | $M_2: 1.52$ | Nf.Kcl4 | 2.57 | $M_2: 2.16$ |
| Ywt.Kcl4 | 1.68 | | Wt.Kcl5 | 1.22 | $s^2_1 = 0.56$ | Nf.Kcl5 | 2.68 | $s^2_2 = 0.13$ |
| Ywt.Kcl5 | 2.24 | $M_2: 1.98$ | Wt.Kcl6 | 2.46 | | Nf.Kcl6 | 1.85 | |
| Ywt.Kcl6 | 2.2 | $s^2_2 = 0.19$ | Wt.Kcl7 | 2 | | Nf.Kcl7 | 1.76 | |
| Ywt.Kcl7 | 2.3 |  |  |  |  |  |  |  |
| Ywt.Kcl8 | 2.69 |  |  |  |  |  |  |  |

DC = FBM; Kcl = FBM + Kcl (0.4M)

### Additional File 5

**Table S1.**
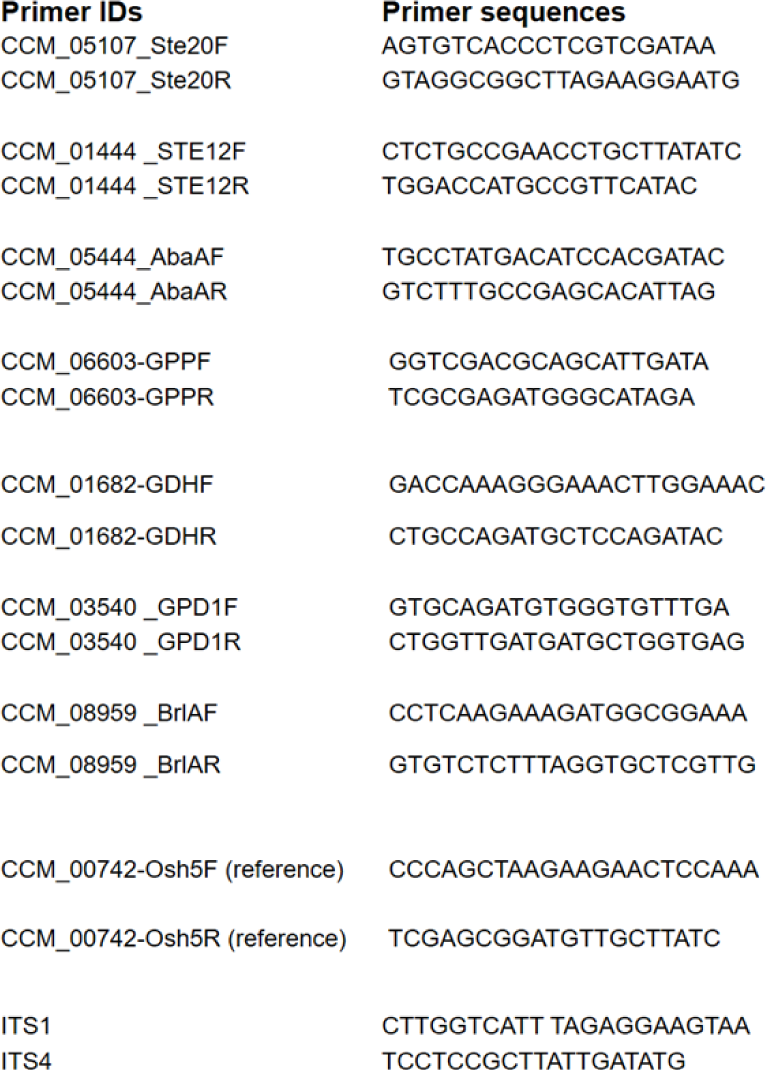
RT-PCR and ITS primer sequences.

**Figure S1.**
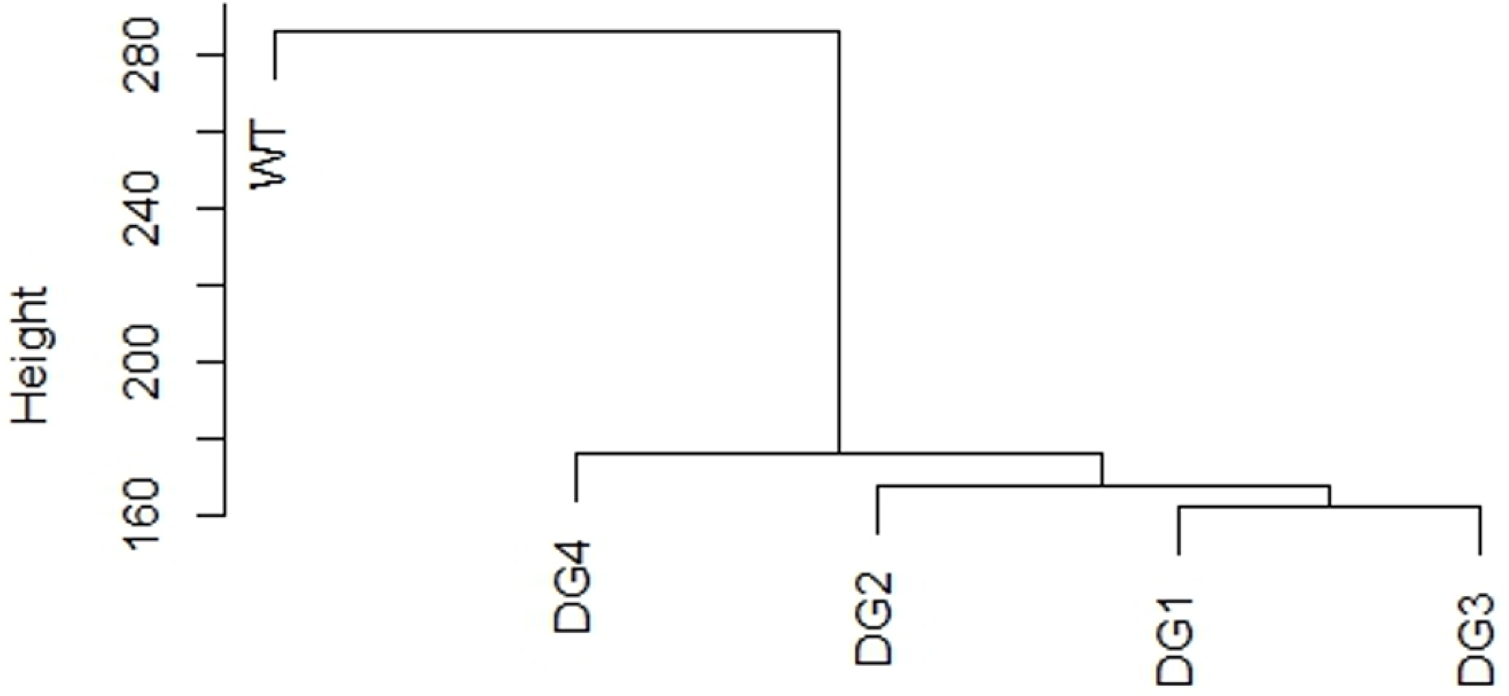
Cluster dendrogram of RNAseq data.

